# Deletion of NuRD component *Mta2* in nephron progenitor cells causes developmentally programmed FSGS

**DOI:** 10.1101/2023.10.18.562984

**Authors:** Jeannine Basta, Lynn Robbins, Lisa Stout, Michelle Brennan, John Shapiro, Mary Chen, Darcy Denner, Angel Baldan, Nidia Messias, Sethu Madhavan, Samir V. Parikh, Michael Rauchman

## Abstract

Low nephron endowment at birth is a risk factor for chronic kidney disease. The prevalence of this condition is increasing due to higher survival rates of preterm infants and children with multi-organ birth defect syndromes that affect the kidney and urinary tract. We created a mouse model of congenital low nephron number due to deletion of *Mta2* in nephron progenitor cells. *Mta2* is a core component of the Nucleosome Remodeling and Deacetylase (NuRD) chromatin remodeling complex. These mice developed albuminuria at 4 weeks of age followed by focal segmental glomerulosclerosis (FSGS) at 8 weeks, with progressive kidney injury and fibrosis. Our studies reveal that altered mitochondrial metabolism in the post-natal period leads to accumulation of neutral lipids in glomeruli at 4 weeks of age followed by reduced mitochondrial oxygen consumption. We found that NuRD cooperated with Zbtb7a/7b to regulate a large number of metabolic genes required for fatty acid oxidation and oxidative phosphorylation. Analysis of human kidney tissue also supported a role for reduced mitochondrial lipid metabolism and ZBTB7A/7B in FSGS and CKD. We propose that an inability to meet the physiological and metabolic demands of post-natal somatic growth of the kidney promotes the transition to CKD in the setting of glomerular hypertrophy due to low nephron endowment.

## INTRODUCTION

Conditions that lead to low nephron endowment, such as congenital renal hypoplasia, intrauterine growth retardation (IUGR) and premature birth are risk factors for later development of chronic kidney disease (CKD) and hypertension (HTN). Clinically, affected individuals develop secondary or adaptive focal segmental glomerulosclerosis (FGSG) manifested by proteinuria and progressive CKD. Congenital anomalies of the kidney and urinary tract (CAKUT) associated with reduced nephron number occur with a frequency of 4-60 in 10,000 live births (1). Preterm births (before 37 weeks gestation) occur at a rate of ∼11% worldwide resulting in 15 million premature births annually (2–4). Globally, IUGR affects ∼10-15% of all pregnancies and the frequency is close to 20% in resource-limited countries (5, 6). In the last 20-30 years, survival of 25-27 weeks old premature newborns has increased substantially in resource rich societies (7, 8). Increased survival of premature infants with birth weights < 1000g is associated with development of CKD starting in childhood and adolescence [reviewed in (9, 10)]. The relatively high prevalence of these clinical conditions makes them significant contributors to the burden of kidney disease. Importantly, while moderate (30-40%) reductions in nephron endowment increase the risk of CKD and HTN, they often go clinically undetected (11, 12).

More than three decades ago, Brenner and colleagues proposed that deficits in nephron number lead to single nephron hyperfiltration and glomerular hypertrophy, ultimately leading to glomerulosclerosis (13). These groundbreaking studies established the pathogenic significance of glomerular hemodynamics in kidney disease. However, the molecular mechanisms that determine whether hypertrophy will become maladaptive, leading to fibrosis and CKD are not well understood. Progress in this field has been hindered by the limitations in existing animal models. Severe acquired reductions in nephrons, as typified by the commonly used 5/6 nephrectomy model in rodents, do not recapitulate congenital defects in nephron formation. During nephrogenesis, genetic and environmental factors that lead to quantitative reductions in nephrons also have the potential to affect proper differentiation of the nephron. Consequently, these qualitative defects, together with the quantitative deficit, likely affect the risk for subsequent development of kidney disease. The metabolic demands of post-natal somatic growth may also promote transition from adaptive to maladaptive hypertrophy when nephron number or differentiation has been impaired during embryonic formation of the kidney.

In this study, we describe a novel genetic model of developmentally programmed CKD due to deletion of *Mta2* during formation of the kidney. *Mta2* is a component of the nucleosome remodeling and deacetylase (NuRD) chromatin remodeling complex. These mutants develop proteinuria by 4-6 weeks and at 8 weeks exhibit histological evidence of CKD with focal segmental glomerulosclerosis, tubulointerstitial fibrosis and inflammatory cell infiltrate. Our results indicate that a moderate reduction of nephron endowment combined with impaired maturation of the nephron leads to CKD. We demonstrate that NuRD is required for regulation of a large number of genes required for mitochondrial metabolism via its interaction with the transcription factor Zbtb7a/b. Overall, this work provides novel mechanistic insights into developmentally programmed CKD and suggests that a failure to meet the physiological and metabolic demands of post-natal growth promote a transition from adaptive to maladaptive hypertrophy in the kidney.

## RESULTS

### Deletion of Mta2 in nephron progenitor cells results in renal hypoplasia

To investigate the role of *Mta2* in nephron progenitor cells, we crossed *Mta2* conditional mutant mice (*Mta2flx/flx*) with *Six2*-Cre. Mta2 protein was not detected in the cap mesenchyme of *Six2*- Cre, *Mta2flx/flx* kidneys, while its expression was preserved in the stroma and ureter (Fig. 1A). In *Mta2flx/flx* control kidneys, Mta2 was detected in nephron progenitor cells, pre-tubular aggregates, comma and S-bodies, and podocytes in developing glomeruli (Fig. 1A). Mta2 was efficiently deleted from these differentiating structures, which are derived from Six2+ progenitor cells, in *Mta2* mutant mice (Fig. 1A). Mutant kidneys appeared grossly normal from embryonic day (E) 11.5-E14.5. Beginning at E15-E16, some *Mta2* mutant kidneys were smaller than wild type littermates, and moderate hypoplasia was apparent in the most affected *Mta2* mutant kidneys (Fig. 1B). Histological examination revealed grossly normal kidneys at E16 with a comparable nephrogenic zone to control littermates (Fig. 1C).

**Figure 1.**
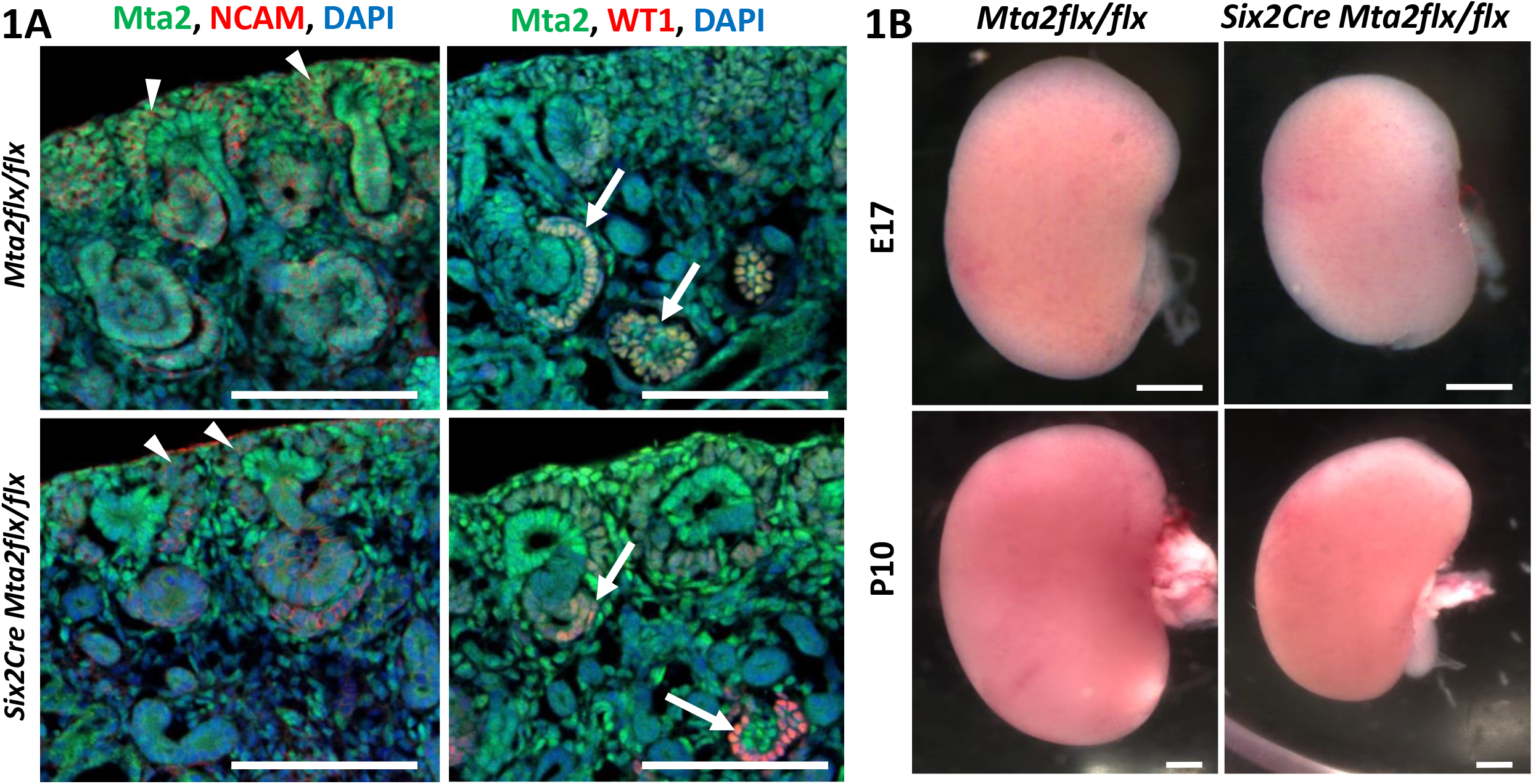
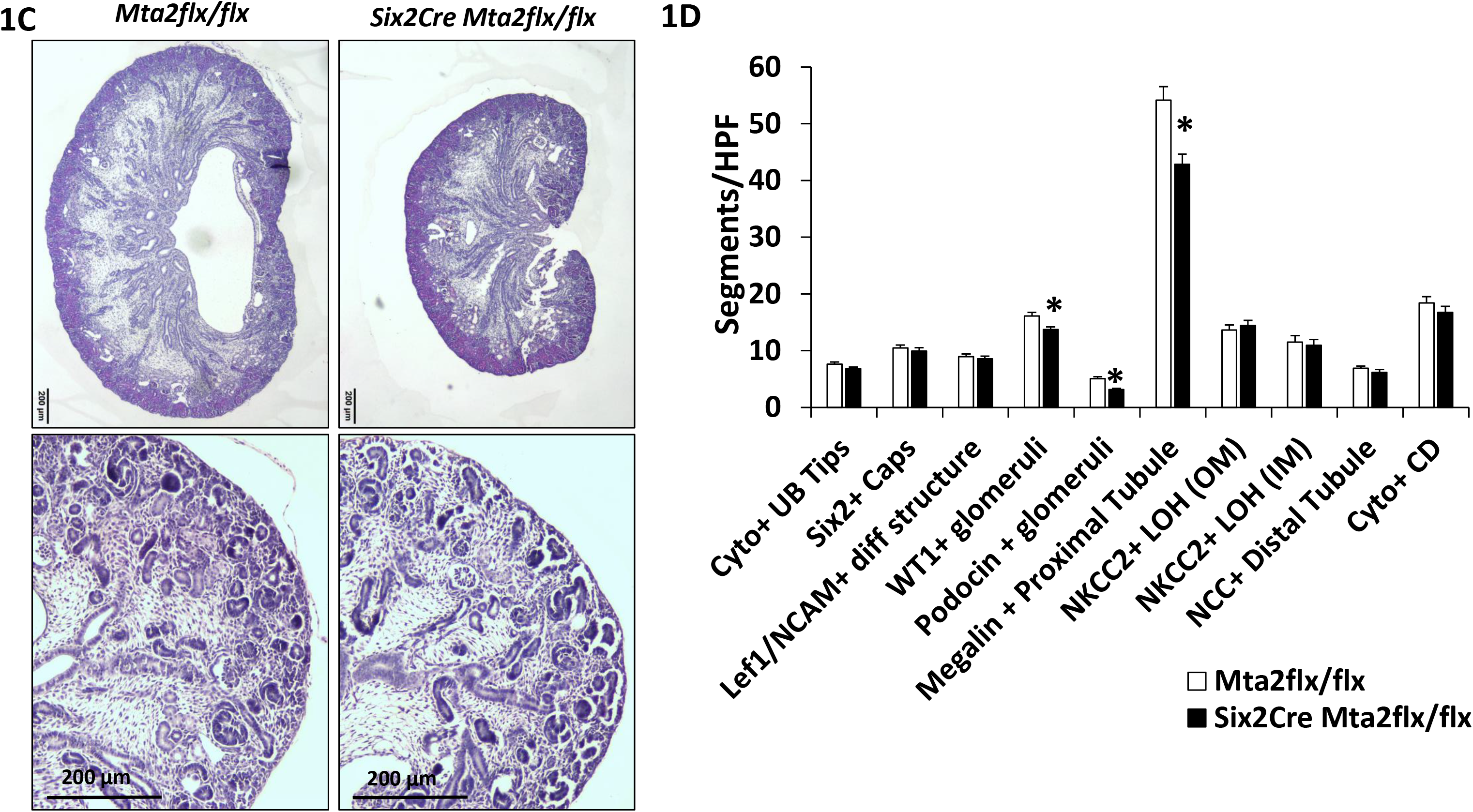
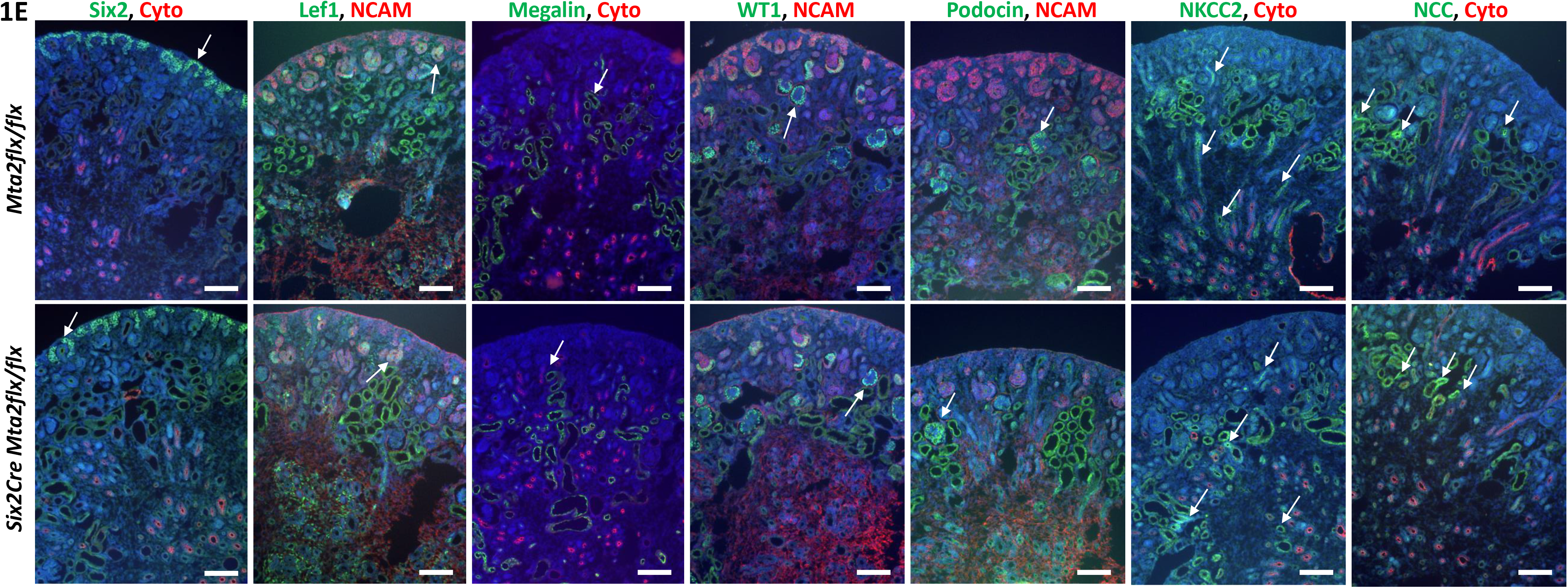
Deletion of *Mta2* in nephron progenitor cells results in renal hypoplasia and impaired nephron formation. **A.** E17 kidney stained for Mta2 (green), NCAM (red, left panels) or WT1 (red, right panels) and DAPI (blue). In control *Mta2flx/flx* kidney Mta2 is expressed ubiquitously including nephron progenitor caps (arrow heads), differentiating nephron structures, and Wt1 positive podocytes (arrows). In mutant *Six2Cre Mta2flx/flx* kidney, Mta2 expression is deleted from nephron progenitor caps (arrow heads), NCAM differentiating nephron structures (comma and S-shaped bodies) and Wt1 positive podocytes (arrows). Scale bar= 100 µm. **B.** Renal hypoplasia is evident in the mutant starting at E15-16. Pictured are representative images from E17 and post-natal day 10 (P10) kidney. E17, scale bar =500 µm. P10 scale bar =750 µm. **C.** Hematoxylin and eosin (H&E) staining of E16 control and mutant kidney. Although mutant kidneys are smaller, a normal nephrogenic zone is present with differentiating nephrons. Scale bar = 200 µm. **D.** Quantification of nephron segments per high powered field (HPF) in E17 kidney using immunofluorescence to identify nephron segments (images in E). For each comparison *Mta2flx/flx* n=3, *Six2Cre Mta2flx/flx* n=3. At least 20 images from at least 12 non-sequential sections for each antibody were quantified. Two-tailed Student’s t-test was used for each antibody comparison between control and mutant. **E.** Immunofluorescence staining in E17 kidney for Six2 positive progenitor cell caps, cytokeratin positive ureteric bud branches, Lef1/NCAM positive differentiating structures (RV, comma body, S-shaped body), Megalin positive proximal tubule, Wt1 positive glomeruli, podocin positive glomeruli, NKCC2 positive loop of Henle, and NCC distal tubule. Arrows indicate examples that were quantified. Scale bar = 100 µm.

### Nephron formation is impaired in Mta2 mutant kidneys

To investigate the etiology of renal hypoplasia, we performed quantitative analyses of ureteric bud (UB) branching, nephron progenitor cells and differentiating structures at E17.5. The number of UB tips per high powered field (HPF) was not reduced in *Mta2* mutants compared with *Mta2flx/flx* controls, indicating that defective UB branching did not account for the phenotype (7.6 ± 0.40 vs. 6.8 ± 0.31, p=0.12) [Fig. 1D, E]. Similarly, the number of Six2 positive caps per HPF (10.48 ± 0.50 vs. 9.92 ± 0.58, p=0.52) were not diminished in the mutant, indicating that a deficiency in nephron progenitor cells did not account for renal hypoplasia in the mutant. Consistent with this observation, cell cycle analysis of FACS sorted Six2-GFP positive cells did not reveal differences in the fraction of cells in G1, S or G2-M between control heterozygotes and mutants at E15.5 and E18.5 (Fig. 2C, D). Phospho-histone H3 (pHH3) staining in E17.5 kidneys did not reveal a difference in the total pHH3 cells/HPF or double positive Six2/pHH3+ nephron progenitor cells in the mutant compared with control kidneys (Fig. 2A, B). We also did not find an increase in apoptosis in Six2 positive nephron progenitor cells in mutants at E17.5 (0.6 ± 0.30 vs 1.0 ± 0.27, p=0.35) or Wt1+ podocytes (0.9 ± 0.286 vs 1.2 ± 0.52, p=0.59) [Fig. 2E, F]. These results indicate that renal hypoplasia in *Mta2* mutants was not due to an impaired branching morphogenesis of the ureter or an inability to expand the nephron progenitor population during formation of the kidney. We also did not observe loss of differentiated podocytes or proximal tubules due to apoptosis in young mice after kidney development was completed [P10 Wt1+ podocytes (0.74 ± 0.22 vs 0.91 ± 0.25 p=0.0.59) and LTL+ proximal tubule cells (5.35 + 0.98 vs 6.42 + 0.81, p=0.40) compared with control kidneys (Fig. 2G, H)].

**Figure 2.**
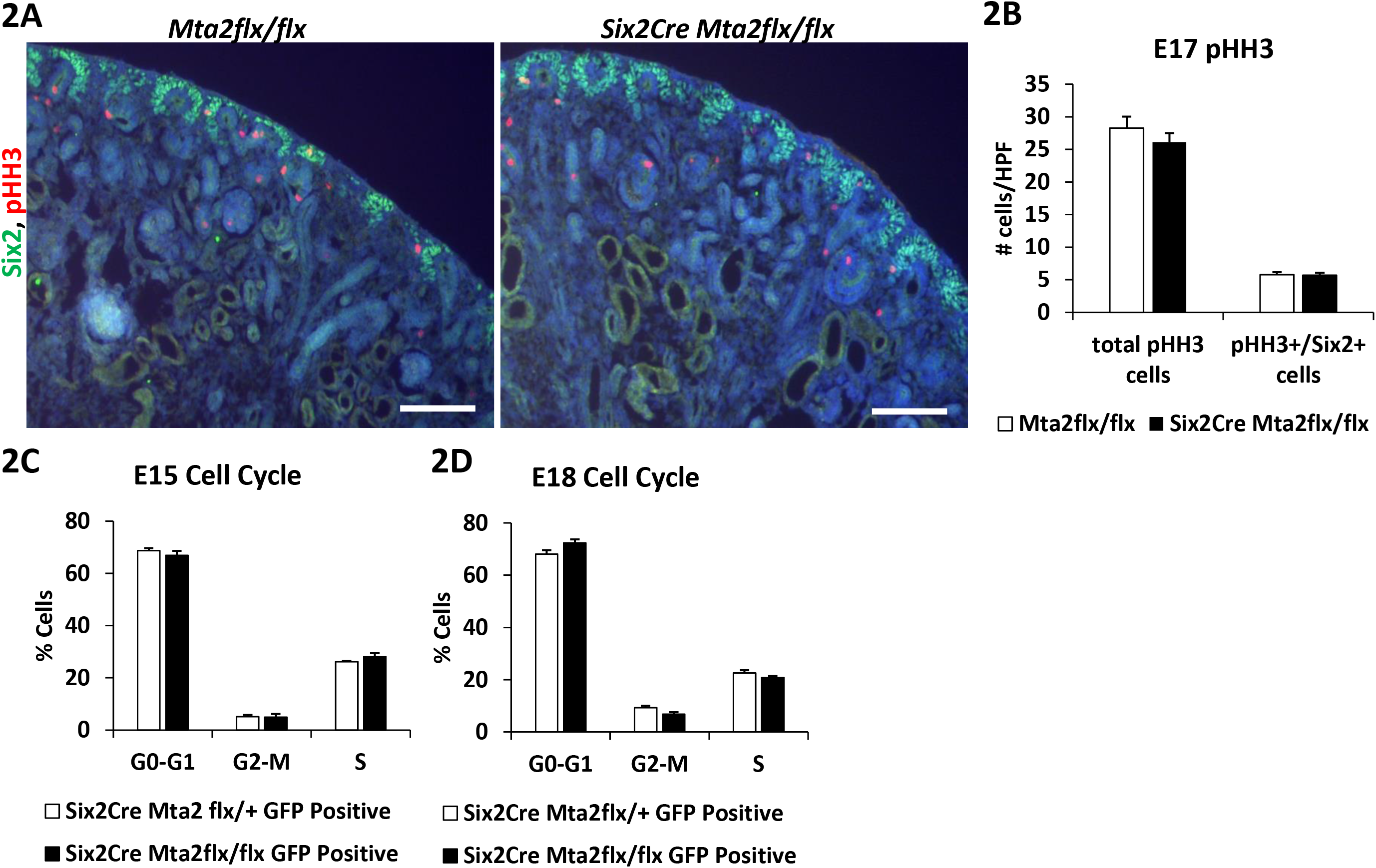
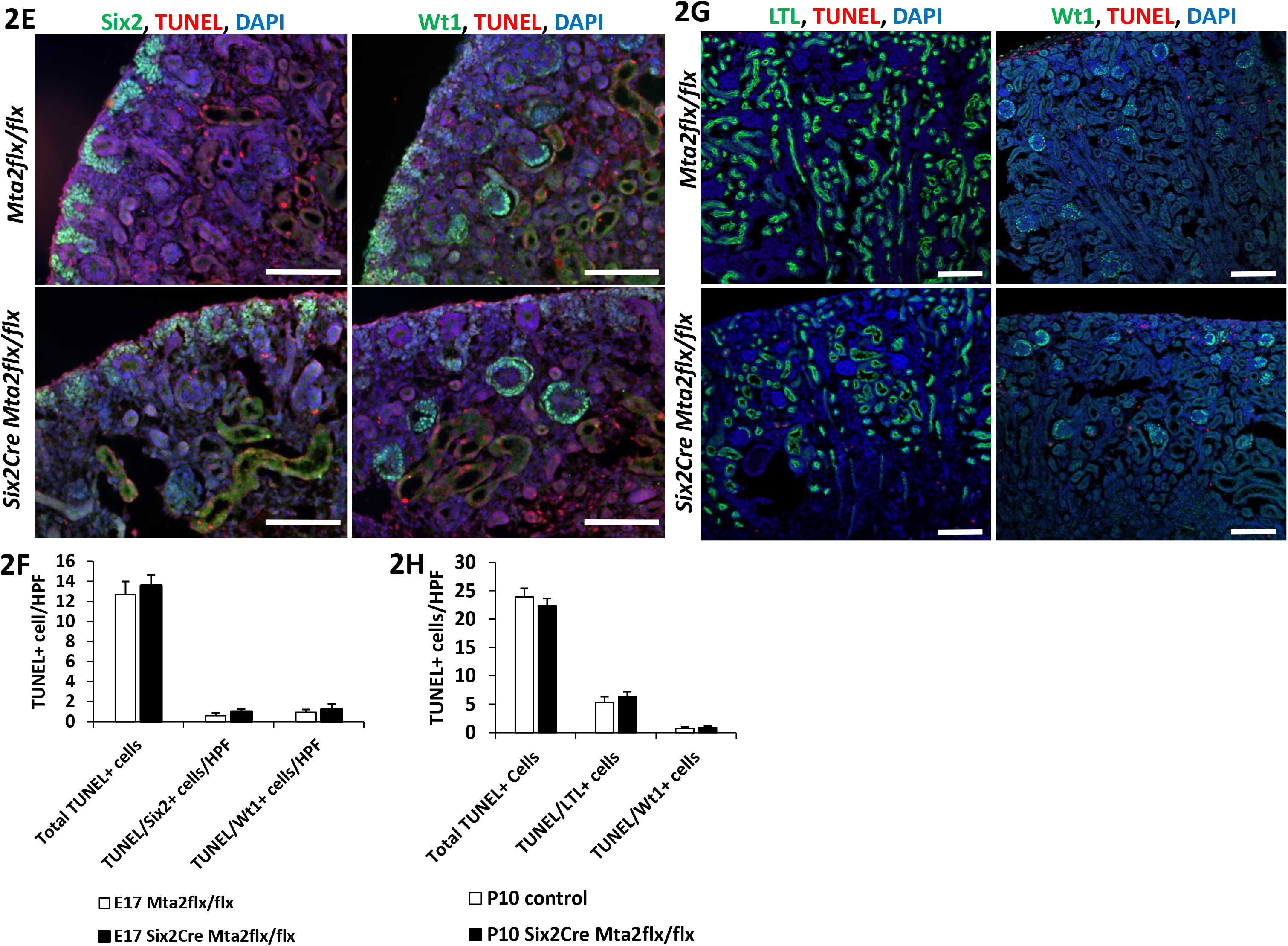
Cell cycle and apoptosis is not changed in *Mta2* mutant embryonic kidney. **A.** Immunofluorescence staining for Six2 positive progenitor cells and pHH3 positive cells in E17 control and mutant kidney. Scale bar = 100 µm. **B.** Quantification of the number of total pHH3 positive cells per HPF and pHH3/Six2 double positive cells per HPF. *Mta2flx/flx* n=3, *Six2Cre Mta2flx/flx* n=3. At least 36 images from at least 12 non-sequential sections were quantified. **C.** Cell cycle analysis performed on E15 and E18 (**D**) Six2 positive progenitor cells from control *Six2Cre Mta2flx/+* and mutant *Six2Cre Mta2flx/flx* kidney. E15 *Six2Cre Mta2flx/+* n=8, *Six2Cre Mta2flx/flx* n=5. E18 *Six2Cre Mta2flx/+* n=7, *Six2Cre Mta2flx/flx* n=2. Two-tailed Student’s t-test was performed between control and mutant for each cell cycle stage. **E.** E17 control and mutant mouse kidney stained for Six2 positive progenitor cells (green, left panels) or Wt1 positive podocytes (right panels), TUNEL (red), and DAPI (blue). *Mta2flx/flx* n=3, *Six2Cre Mta2flx/flx* n=3, at least 24 images from 12 non-sequential slides quantified for each antibody. Scale bar = 50 µm. **F.** Quantification of total TUNEL positive cells per HPF, total TUNEL/Six2 double positive cells per HPF, and total TUNEL/Wt1 double positive cells per HPF. E17 *Mta2flx/flx* n=3, *Six2Cre Mta2flx/flx* n=3. Quantification was performed using at least 24 images from 12 non- sequential sections for each antibody combination. **G.** P10 control and mutant kidney stained for LTL (green, left panels) or Wt1 (green, right panels), TUNEL (red), and DAPI (blue). Scale bar = 100 µm. **H.** Quantification of total TUNEL positive cells per HPF, total TUNEL/LTL double positive cells per HPF, and total TUNEL/Wt1 double positive cells per HPF. *Mta2flx/flx* n=3, *Six2Cre Mta2flx/flx* n=3, quantification was performed on at least 22 images from 12 non- sequential sections for each antibody.

To determine if nephron formation was affected in mutants, we quantitated renal vesicles and differentiating nephron structures. We found that Lef1/NCAM positive early differentiating structures were not reduced in *Mta2* mutant kidneys at E17.5, indicating that nephron induction was not impaired in the mutant (Fig. 1D, E). However, quantitative analysis of differentiating nephrons at E17 revealed a 39% reduction in Podocin+ glomeruli (5.06 ± 0.33 vs. 3.12 ± 0.27, p=5.0 x 10^-6^), a 15% reduction in developing and fully formed Wt1+ glomeruli (16.1 ± 0.65 vs. 13.72 ± 0.46, p=0.003), and a 21% reduction in Megalin+ proximal tubules (54.16 ± 2.36 vs. 42.87 ± 1.77, p=0.0003) [Fig. 1D, E].

To gain insight into the gene expression program regulated by *Mta2* in the kidney, we performed RNA sequencing (RNA-seq) at E17.5 on *Mta2* mutant and developmental stage matched control kidneys (Supp. Table 1). Transcriptional profiling resulted in 247 upregulated and 206 downregulated genes (>1.5 fold, p<0.05) [Fig. 3A]. Gene ontology analysis (Fig. 3B) revealed significant enrichment for multiple metabolic pathways in the downregulated gene set, with many related to lipid metabolism. RNA polymerase II dependent transcriptional activity was also enriched, consistent with the broad role of the NuRD complex regulating gene expression by modifying chromatin. We compared the global gene expression changes in *Mta2* mutants to the top 20 genes expressed in specific cell clusters in mouse embryonic kidney determined by single cell RNA-seq (scRNA-seq) [Supp. Fig. 1, (14)]. We found that most genes that are early differentiation markers of specific nephron segments by scRNA-seq were downregulated in the *Mta2* mutant. We also found that terminal differentiation genes in podocytes (*Mafb*, *Nphs1*, *Nphs2*), and renal tubular epithelia genes encoding transporter and ion channel proteins, were downregulated in the mutant. These results indicate that nephron formation and their proper terminal differentiation were impaired due to deletion of *Mta2* in Six2+ nephron progenitor cells.

**Figure 3.**
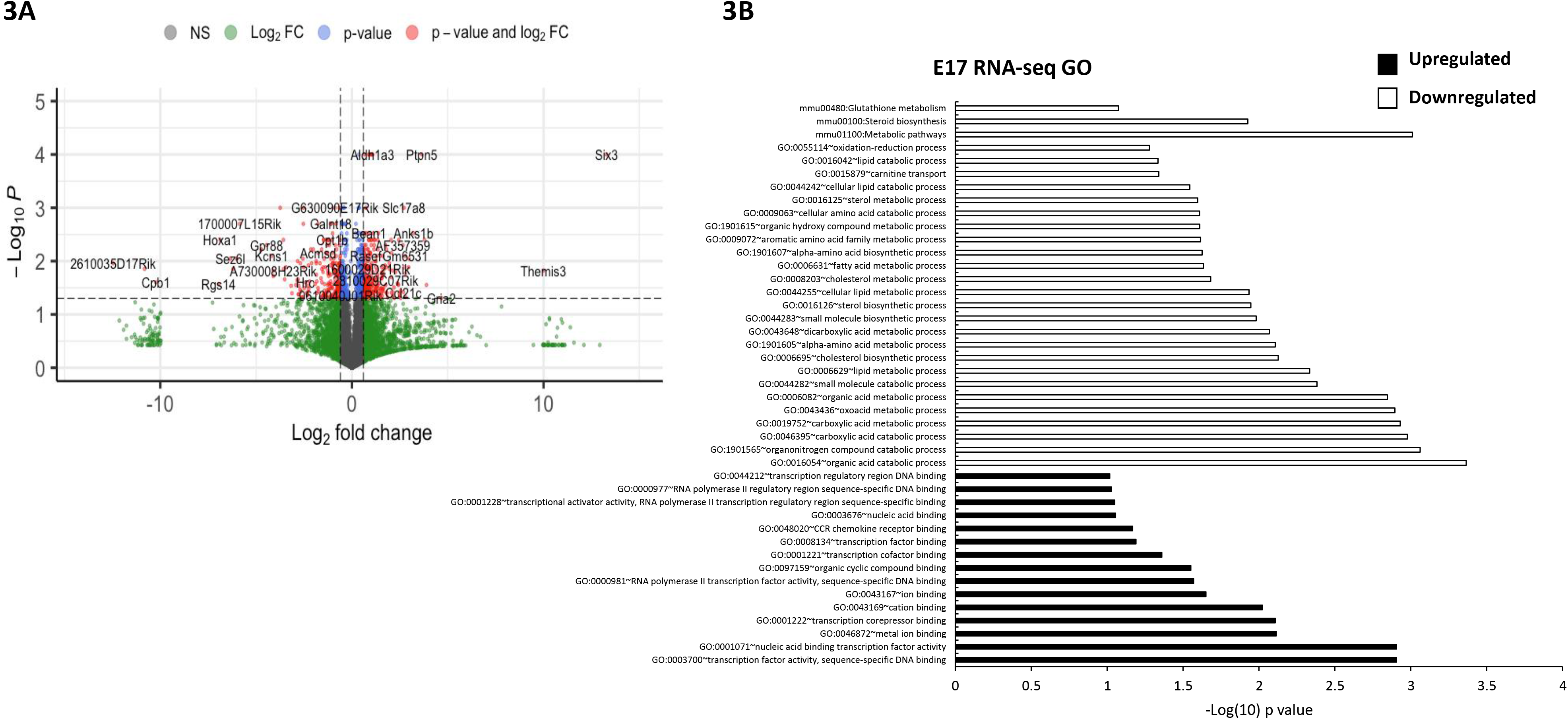
E17 RNA-seq on control and mutant *Mta2* kidney. **A.** Volcano plot of E17 RNA-seq of mutant kidney depicting genes with 1.5 fold change, p < 0.05 (red). **B.** Gene ontology analysis of upregulated and downregulated genes. Downregulated genes are enriched for many metabolic processes including lipid metabolism and fatty acid metabolism. Upregulated genes are enriched for transcriptional regulation gene ontologies including transcription factor cofactor binding, nucleic acid binding, and transcription regulatory region binding.

### Mta2 mutant mice are viable and develop focal segmental glomerulosclerosis (FSGS)

*Mta2* mutant mice were viable and fertile, and appeared healthy. Body weights in male and female mice at post-natal time points (P0-9 months) were not significantly different (Supp. Fig. 2). However, at 2 months of age, male and female *Mta2* mutant mice exhibited significant albuminuria (1416 ± 295.1 µg albumin/mg creatinine), which was not present in *Mta2flx/flx* control mice (36.11 ± 4.4 µg albumin/mg creatinine) [Fig. 4A]. Periodic Schiff (PAS) staining of kidneys revealed histological lesions characteristic of focal segmental glomerulosclerosis (FSGS), with segmental areas of PAS positive material in mutant but not control glomeruli (Fig. 4C). At 6 months of age, there was progression of the histological lesions, which included globally sclerotic glomeruli, increased mesangial matrix and hypercellularity, and proliferation of parietal epithelial cells leading to pseudo-crescent formation (Fig. 4C, Supp. Fig. 3). Survival of *Mta2* mutant mice was significantly decreased after 6 months of age (Fig. 4B), which corresponded to the presence of severe kidney injury and fibrosis. Progression of FSGS lesions from 2 to 6 months was confirmed by worsening of the glomerulosclerosis score (Fig. 4D, Supp. Fig. 3). Trichrome staining confirmed the presence of segmental and global scarring of glomeruli, and revealed the development of interstitial fibrosis in 6 months old mutant kidneys (Supp. Fig. 4). There was also evidence of tubule injury and interstitial inflammation, with progressive worsening of tubule-interstitial injury score from 2 to 6 months (Fig. 4E, Supp. Fig. 5, 6). Immunofluorescence staining for T cells (CD3) and macrophages (F4/80) showed significant increases of these immune cells in interstitial spaces of mutant kidney at both 2 and 6 months of age (Fig. 4F). We performed TUNEL staining to determine if cell death due to apoptosis was contributing to the observed phenotype. We did not detect an increase in TUNEL+ cells in Wt1+ podocytes at 2 months of age, when FSGS is first clinically apparent in these mice. However, TUNEL+ cells were markedly increased in LTL+ proximal tubule epithelia and interstitial cells at 2 months of age, indicating ongoing tubulo-interstitial injury (Fig. 4G, H).

**Figure 4.**
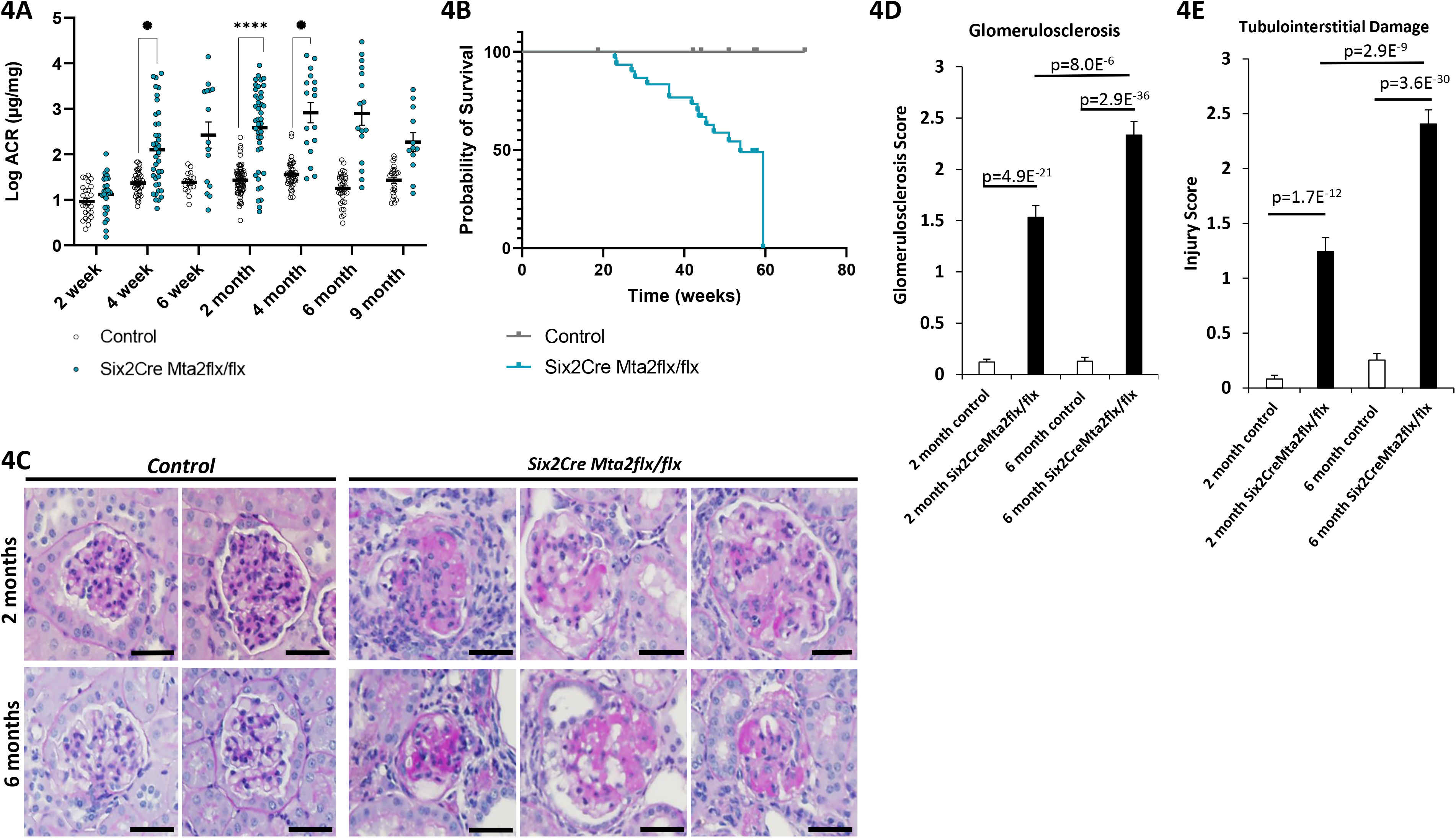
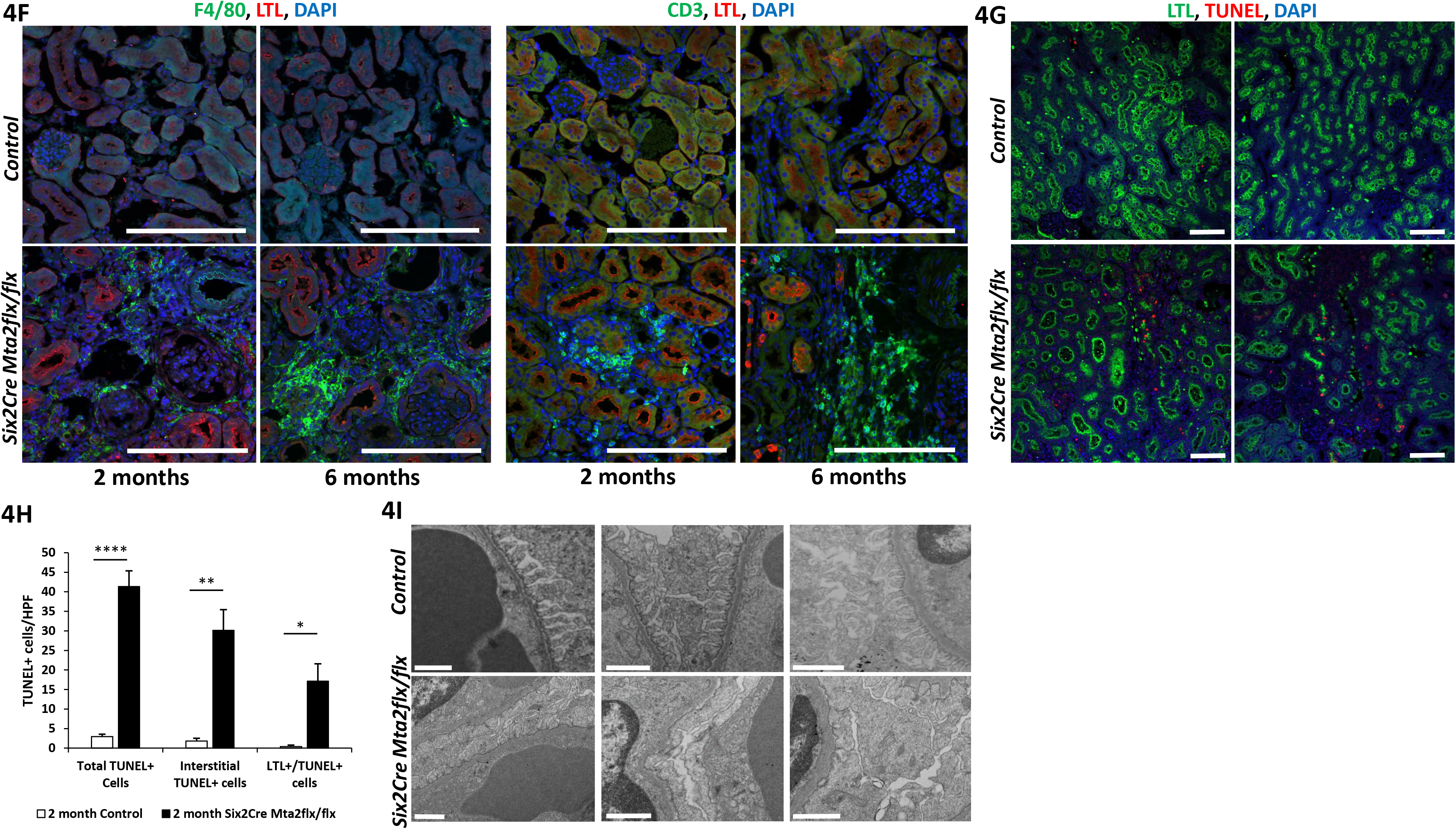
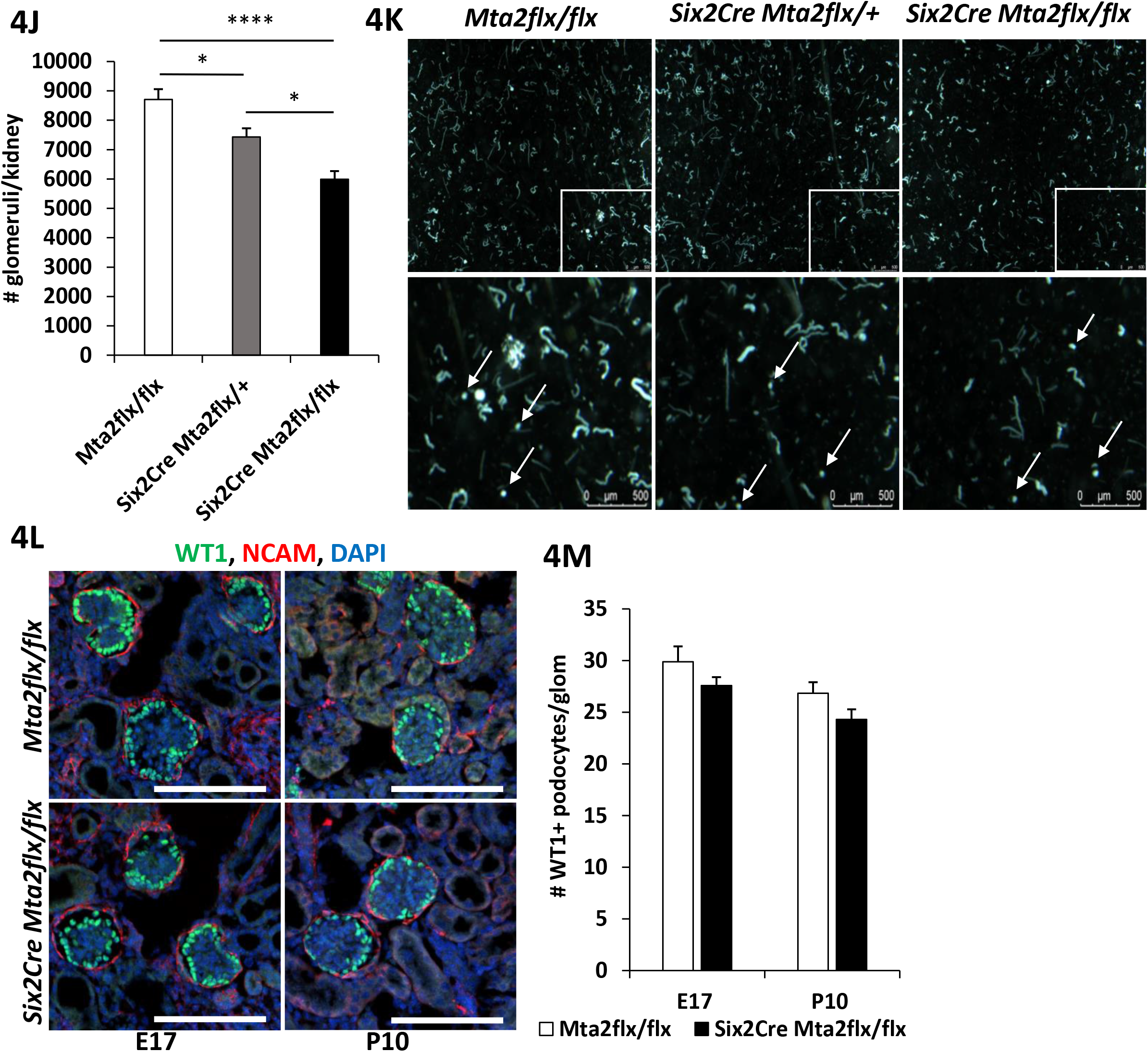
*Mta2* mutant mice develop FSGS. **A.** Urine albumin normalized to urine creatinine, displayed as the log of the albumin to creatinine ratio (Log ACR), µg/mg, measured in control and mutant mice 2 weeks to 9 months of age. Statistics analyzed using multiple paired t-tests (Prism). 4 weeks p=0.02, 2 months p = 0.000026, 4 months p= 0.045. **B.** Kaplan-Meier survival curve of control and mutant mice. Statistical analysis using Log-rank (Mantel-Cox test), p= 0.0034. Hazard Ratio (Mantel-Haenszel) 4.893 (95% Cl, 1.692 - 14.15) (Prism). Control n=15 (male n=6, female n=8), *Six2Cre Mta2flx/flx* n= 30 (male n= 16, female n= 14). **C.** Periodic acid- Schiff (PAS) stained sections of control and mutant kidney at 2 months and 6 months of age. Focal segmental lesions present at both 2 months and 6 months of age. Scale bar = 50 µm. **D.** Semi-quantitative analysis of glomerulosclerosis (grade 0, no change – grade 4, >75% of glomerulus PAS positive). At least 40 glomeruli were evaluated from 3 biological replicates for each genotype. Results are the mean ± SEM. Statistical analysis was performed using a two- tailed Student’s t-test. Representative images that were analyzed are shown in **Supp.** Fig. 3. **E.** Semi-quantitative analysis of tubulointerstitial damage evaluated by determining the % damage per HPF (grade 0, no change – grade 4 >75%). At least 16 individual PAS stained images (**Supp.** Fig. 6) were evaluated from 3 biological replicates for each genotype. Results are the mean ± SEM. Statistical analysis was performed using a two-tailed Student’s t-test. **F.** Immunofluorescence of 2 and 6 month kidney sections for F4/80 (green, left panels) or CD3 (green, right panels), LTL (red) and DAPI (blue). F4/80 macrophages and CD3 positive T-cells were present in mutant kidney at 2 and 6 months of age. Scale bar = 100 µm. **G.** TUNEL (red), LTL (green) and DAPI (blue) staining on 2 month kidney sections from control and mutant. Scale bar = 100 µm. **H.** 2 month kidney quantification of total TUNEL positive cells per HPF, TUNEL positive cells within tubulointerstitium per HPF, and TUNEL positive cells in LTL positive cells per HPF. Results are the mean ± SEM. At least 6 images from 3 biological replicates for each genotype were quantified. Statistical analysis performed using a two-tailed Student’s t-test, **** p= 7.22E-11, ** p=0.0012, *p=0.01. Scale bar = 100 µm. **I.** Transmission electron microscopy of control and mutant 2 month kidney. Defined podocyte foot processes are present in the control kidney. Diffuse podocyte foot effacement is present in the mutant kidney. Scale bar = 1 µm. **J.** Quantification of glomeruli using a crude glomeruli prep from P10 kidney, pictured in (**K**). Compared to control *Mta2flx/flx* kidney, heterozygous mutant and homozygous mutant kidneys had statistically significantly less glomeruli/kidney. Results are the mean ± SEM. Statistical analysis performed using a one-way ANOVA and Tukey’s multiple comparisons test, * p= 0.01, **** p< 0.0001. *Mta2flx/flx* n=3, Six2Cre Mta2flx/+ n=3, *Six2Cre Mta2flx/flx* n=2. **K.** Bright field images of crude glomeruli prep with inserts (boxes in upper panels) magnified in lower panels. Arrows point to glomeruli. Scale bar = 500 µm. **L.** Immunofluorescent staining and confocal imaging of Wt1 in podocytes (green), NCAM (red), and DAPI (blue) of E17 and P10 control and mutant kidney. Scale bar = 100 µm. **M.** Quantification of Wt1 positive podocytes per glomeruli in E17 and P10 control and mutant kidney. At least 30 glomeruli for each genotype and stage were quantified from 3 biological replicates. No significant difference detected between control and mutant at either stage using a two-tailed Student’s t-test. Scale bar = 100 µm.

Consistent with the finding of significant albuminuria, we found podocyte foot process effacement on electron microscopy in *Mta2* mutants (Fig. 4I). Podocyte foot process effacement is a key morphological feature associated with proteinuria. The degree of effacement is thought to distinguish primary FSGS, which typically displays severe diffuse effacement, from genetic and adaptive/secondary FSGS in which effacement is segmental or less severe. At 2 months of age, foot process effacement in *Mta2* mutants was segmental, not diffuse, indicative of genetic or adaptive FSGS. Because *Mta2* mutants exhibited renal hypoplasia, we quantitated the reduction in nephron number, which is a risk factor for adaptive glomerulosclerosis (13, 15). We found that the number of glomeruli per kidney was reduced by 31% in *Mta2* mutants compared with *Mta2flx/flx* controls at postnatal day (P)10 (5990 vs. 8708, *Mta2flx/flx* n=3, Six2Cre *Mta2flx/flx* mutant n=2, p=1.0 X 10^-4^), about 4-6 days after nephrogenesis is completed in the mouse (Fig. 4J, K). Glomerular number was also significantly reduced in *Mta2* heterozygotes (15%, p=0.01), but to a lesser extent than in homozygous mutants. In addition to glomerular number, a reduction in the number of podocytes, as a consequence of the improper glomerulogenesis in *Mta2* mutants, could contribute to development of FSGS. However, we did not detect a reduction in the number of Wt1-positive cells per glomerulus during formation of glomeruli at E17.5, nor at P10 after nephrogenesis was completed (Fig. 4L, M), indicating that reduced podocyte formation during development was not a factor in causing FSGS in these mice. These results indicated that reduced nephron endowment contributed to development of proteinuria and FSGS in this mutant.

The temporal relationship between morphological changes in the kidney and the onset of albuminuria in genetic and adaptive FSGS is not clear. To define the earliest manifestations of disease in *Mta2* mutant mice, we quantitated albuminuria and examined kidney histology prior to 2 months of age, when albuminuria was severe and FSGS lesions were apparent. At 4 weeks of age, *Mta2* mutant mice had significant albuminuria (Fig. 4A), though this was less severe and more variable than at 8 weeks. Histological analysis of kidneys in 4-week-old mice displayed no pathological features and these kidneys were indistinguishable from litter mate controls (Supp. Fig. 7A). Congenital nephron deficit is associated with glomerular hypertrophy. At 4 weeks of age, we found that glomeruli in *Mta2* mutant kidneys were significantly larger than control littermates (Supp. Fig. 7B-D). The gradual onset of proteinuria with normal histology, glomerulomegaly and subsequent progression to glomerulosclerosis is consistent with adaptive/secondary FSGS.

### Transcriptional profiling in Mta2 mutant adult kidneys reveals impaired differentiation and metabolic gene expression

While reduced nephron number is a risk factor for glomerulosclerosis, the moderate reduction observed suggested that this factor alone did not likely account for glomerular disease in *Mta2* mutant mice. To explore the molecular mechanism of FSGS in this model, we performed RNA-seq on kidneys from 2 months old mutant and control mice (Supp. Table 2). Transcriptional profiling revealed 210 upregulated and 637 downregulated genes (>1.5 fold, p<0.05) [Fig. 5].

**Figure 5.**
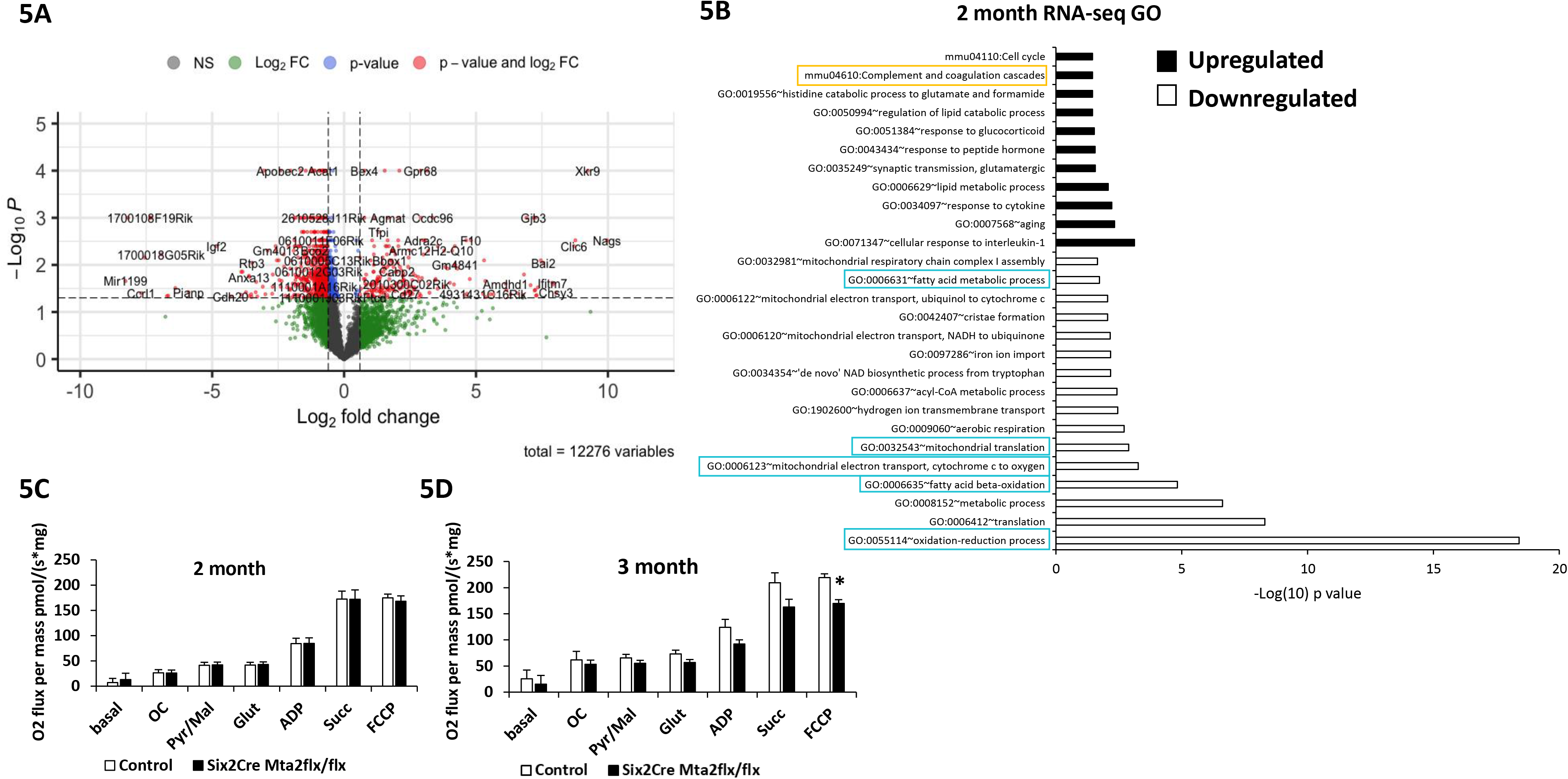
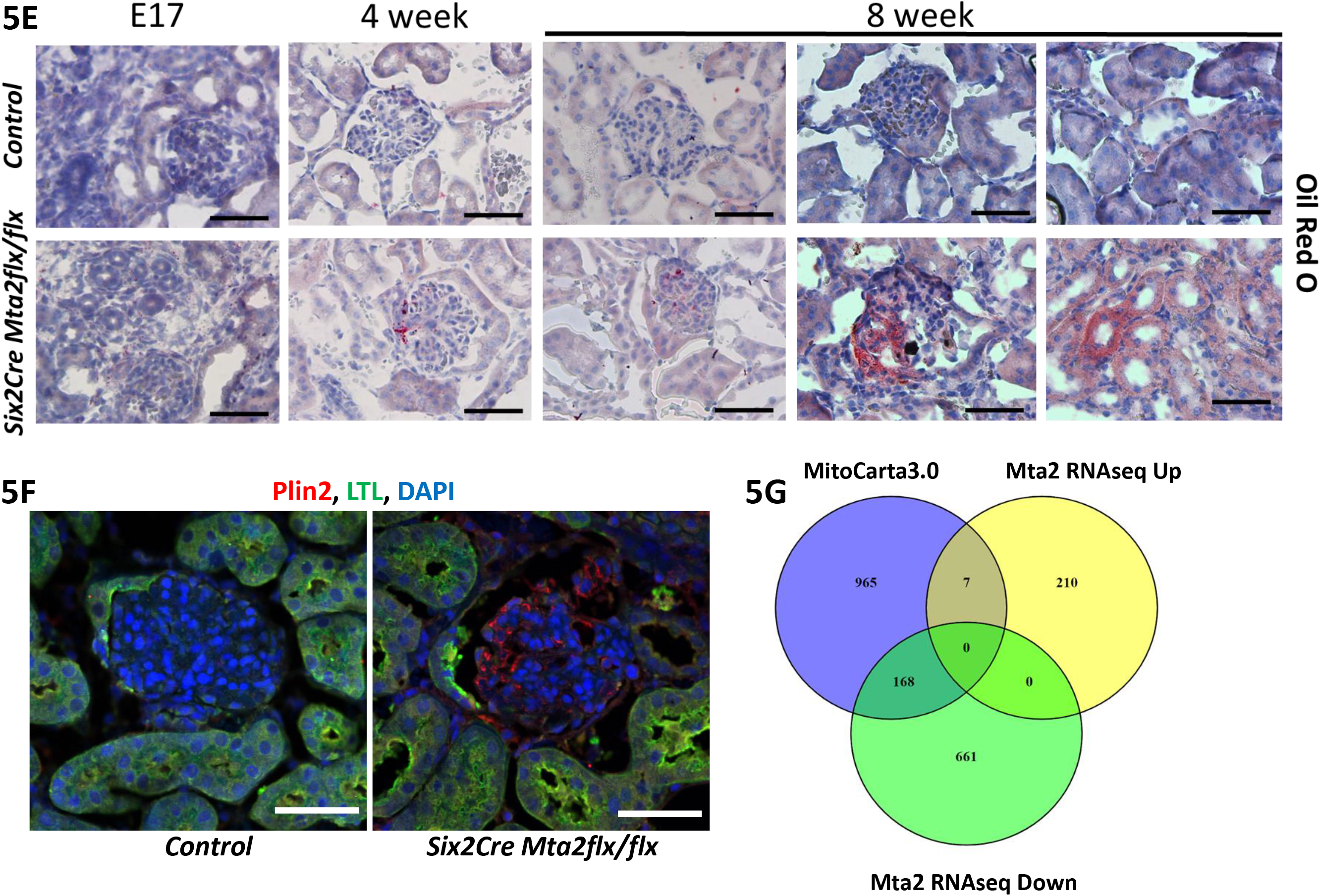
Transcriptional profiling in *Mta2* mutant adult kidney reveals altered metabolic gene expression. **A.** Volcano plot of RNA-seq of 2 month mutant kidney depicting significantly changed genes (red) with 1.5 fold change in expression, p < 0.05. **B.** Gene ontology analysis of upregulated and downregulated genes. Downregulated genes are significantly enriched for mitochondrial pathways (oxidation-reduction, electron transport, complex I assembly) and fatty acid metabolism. Upregulated pathways are enriched for complement and coagulation cascades and response to cytokines. **C.** Oroboros high resolution respirometry performed in control and mutant kidney cortex at 2 and 3 (**D**) months of age. Mitochondrial respiration was measured with OROBOROS Oxygraph system in permeabilized Kidney with sequential additions of octanoyl-L-Carnitine (OC), pyruvate and malate (Pyr/mal), glutamate (G), adenosine diphosphate (ADP), succinate (Succ) and FCCP. Results are represented as the mean ± SEM and were analyzed by multiple paired t tests, p= 1.01E-5 for 3 month FCCP. 2 month control n=11, *Six2Cre Mta2flx/flx* n=11. 3 month control n=10, *Six2Cre Mta2flx/flx* n=11. **E.** Oil red O staining on kidney sections from control and mutant E17, 4 week, and 8 week old mice. A low level ubiquitous increase in oil red O deposition in E17 mutant kidney was observed. In 4 week old mutant kidney oil red O staining was observed in glomeruli, and at 8 weeks increased oil red o staining was observed in glomeruli and tubules. Scale bar = 50 µm. **F.** Immunofluorescence staining in 8 week old control and mutant kidney sections for Plin2 (red), LTL (green), and DAPI (blue). Increased Plin2 staining is observed in glomeruli of mutant kidney. Scale bar = 100 µm. **G.** Venn diagram comparing 2 month *Mta2* RNA-seq differentially expressed genes with the MitoCarta3.0 database of genes. 168 downregulated genes (20%) in the 2 month *Mta2* mutant have expression and function in the mitochondria.

Our analysis at E17.5 indicated that terminal differentiation genes for nephron segments derived from Six2 positive cells were downregulated in the mutant, suggesting that maturation of the nephron was not complete in the mutant. The transcriptional profile at 2 months also revealed that many terminal differentiation genes in podocytes and renal tubular segments were downregulated (Supp. Fig. 8). These data suggest that impaired terminal differentiation of Six2- derived nephron segments persisted in the adult mutant.

Gene Ontology analysis (Fig. 5B) identified oxidation-reduction phosphorylation (Ox Phos) as one of the most highly enriched pathways that was downregulated. Multiple genes from Ox Phos complexes I-IV exhibited reduced expression. Similar to E17.5, lipid/fatty acid metabolism genes were also significantly affected. Another highly enriched biological process identified was protein translation, which includes multiple mitochondrial protein translation genes that were downregulated. These results implicate mitochondrial dysfunction in the pathogenesis of FSGS in *Mta2* mutants. Together, these findings suggest that post-natal maturation of the kidney was impaired in the mutant, resulting in an inability to meet the physiological and metabolic demands during growth and in adulthood.

### Lipid accumulation and impaired mitochondrial function in Mta2 mutants

To investigate whether mitochondrial metabolism was affected in *Mta2* mutant kidneys, we performed high-resolution respirometry on fresh kidney cortex. At 3 months of age, maximal respiration was significantly reduced in *Mta2* mutant cortex (Fig. 5D). These findings are consistent with the large number of oxidative phosphorylation (OxPhos) genes that are reduced in the mutant. However, we did not detect a difference in maximal respiration at 2 months of age (Fig. 5C), when FSGS is first apparent, suggesting that while reduced OxPhos may contribute to disease progression it is not likely the initiating event. Our RNA-seq studies revealed that expression of many genes involved in lipid metabolism was altered at both E17.5 and 2 months of age. Altered lipid metabolism can lead to accumulation of lipids in podocytes and tubular epithelia causing mitochondrial dysfunction and cell injury. To examine this possibility, we performed Oil Red O staining in control and *Mta2* mutant kidneys. At 2 months of age, we detected neutral lipid accumulation in glomeruli in mutant but not control kidneys (Fig. 5E). At 4 weeks of age, when proteinuria was first noted but prior to onset of histological FSGS, lipid accumulation was also detected in mutant glomeruli (Fig. 5E). Lipid deposition was confirmed with Plin2 (ubiquitously expressed and located on lipid droplet particle surfaces) antibody staining. Increased Plin2 staining was observed in 2 month mutant glomeruli (Fig. 5F).

Together, these studies indicate that altered lipid metabolism was an early finding in *Mta2* mutants and likely contributed to development of mitochondrial dysfunction detected by respirometry at 3 months of age.

### NuRD and Zbtb7a occupy many common gene promoters of metabolic genes with altered expression in Mta2 mutants

We compared our RNA-seq results to MitoCarta3.0, an inventory of mammalian mitochondrial proteins (16). The overlap of downregulated genes in the *Mta2* mutant (p<0.05) with mitochondrial proteins was 20% (168/829) (Fig. 5G, Supp. Table 3). This analysis also revealed that transcriptional regulators in the mitochondria expression neighborhood included Mta2 (17), suggesting that NuRD may directly regulate expression of genes required for mitochondrial function. To test this possibility, we performed ChIP-seq on kidney cortex at 2 months of age using an Mta2 specific antibody (Supp. Table 4). Mta2 binding was detected in the promoters of many genes that showed reduced expression (<1.5 fold, p < 0.05) in the mutant (443/637, 70%, Fig. 6B, Supp. Table 5). We performed gene set enrichment analysis (GSEA) of these 443 genes using the Hallmark gene sets. This analysis showed significant enrichment for genes involved in oxidative phosphorylation (p-value =1.44E-24, FDR q-value=7.20E-23), metabolism of lipids (Adipogenesis, p-value = 1.98E-07, FDR q-value =4.95E-06) and fatty acid metabolism (p-value=4.28E-05, FDR q-value=4.28E-04) [Supp. Table 5]. To validate that NuRD bound these genes’ promoters, we also performed ChIP-seq for another core NuRD component, Mbd3 (Supp. Table 4). Genome wide in control kidneys, Mbd3 was bound at a high percentage of genomic sites also bound by Mta2 (96% within active promoter regions, 97% within predicted enhancers, 75% within intronic regions, and 80% within exons), consistent in their role together in the NuRD complex. Together, these results indicate that NuRD regulates expression of genes required for mitochondrial function and lipid metabolism, likely via direct binding at regulatory regions in their promoters.

**Figure 6.**
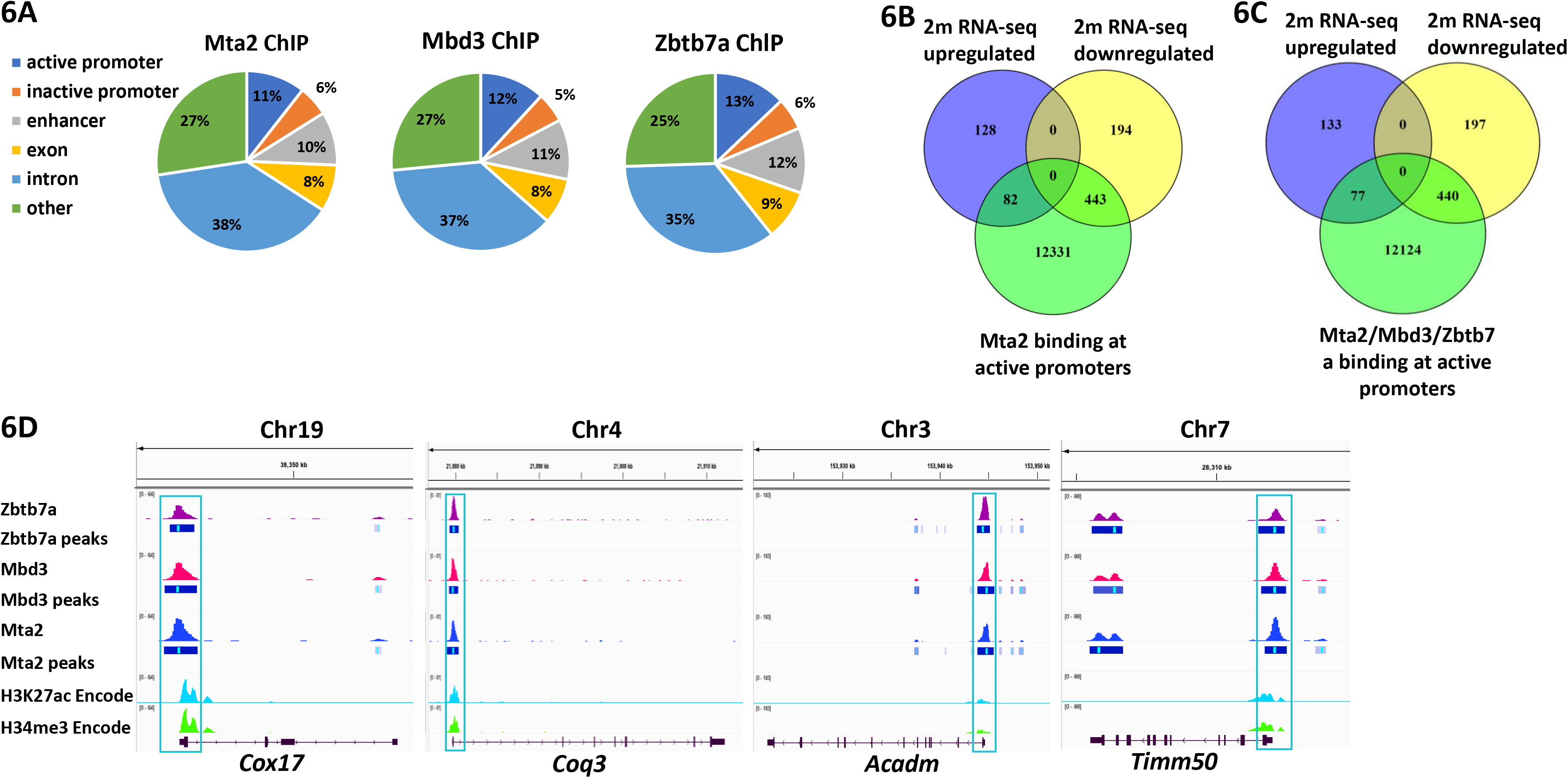
NuRD and Zbtb7a bind common promoters of metabolic genes in *Mta2* mutant kidneys. **A.** Annotation of ChIP-seq binding sites for Mta2, Mbd3, Zbtb7a in kidney cortex. Genome wide, binding patterns for NuRD (Mta2, Mbd3) is very similar to binding of Zbtb7a, with a majority of binding sites within active promoters of genes. **B.** Venn diagram comparing Mta2 ChIP-seq binding in active promoters in kidney cortex with differentially expressed genes in 2 month *Mta2* mutant kidney. **C.** Venn diagram comparing active promoters of genes co-bound with Mta2, Mbd3, and Zbtb7a and differentially expressed genes in 2 month *Mta2* mutant kidney. **D.** ChIP-seq tracks of Mta2, Mbd3, and Zbtb7a binding in 2 month old kidney cortex with called peaks. ENCODE ChIP-seq of two month old mouse kidney for H3K27ac (ENCFF312MNA) and H3K4me3 (ENCFF673XLI). Promoter region (blue box) of *Cox17* (cytochrome C oxidase, functions in the transfer of electrons from cytochrome C to oxygen), *Coq3* (coenzyme Q, critical component of electron transport chain), *Acadm* (medium chain acyl-CoA, required for fatty acid oxidation), *Timm50* (inner mitochondrial membrane translocase, responsible for maintaining membrane permeability). *Cox17* scale 0-64, *Coq3* scale 0-61, *Acadm* scale 0-183, *Timm50* scale 0-88).

The actions of chromatin remodeling complexes such as NuRD must be coordinated with sequence specific transcription factors to regulate gene expression in specific cells and in response to physiological stresses or injury. We performed gene set enrichment analysis to identify transcription factor binding sites within promoter regions of sets of genes. Using the 443 genes that had decreased expression by RNA-seq in *Mta2* mutant kidneys and Mta2 binding within the promoter region of the gene, we found that Zbtb7a/b was among the top ranked transcription factor binding sites within these promoters (p=5.29E-15, FDR q-value =3.67E-12) [Supp. Table 6]. A consensus DNA motif for Zbtb7/ab was found in 62/443 (14%) of genes with significantly reduced expression, which included many nuclear encoded genes that function in the mitochondria (*Nubpl*, *Cox7c, Cox17, Cox8a*, *Timm50*, *Slc25a19, Mrps31, Mrps28, Ears2*). Zbtb7a and 7b interact with NuRD via their BTB domains to regulate gene expression in hematopoietic and immune cells (18). *Zbtb7b* null mice have impaired adipocyte differentiation and exhibit lipid accumulation (19). We hypothesized that Zbtb7a/b cooperates with NuRD to regulate metabolic genes in the kidney. ChIP-seq for Zbtb7a (Supp. Table 4) revealed that Zbtb7a bound a high percentage of binding sites also bound by Mta2 and Mbd3 (>97% within active promoters, 96% within predicted enhancers, 79% within intronic regions, and 87% within exons). Of the 637 down regulated genes (<1.5fold, p<0.05) in the 2 month *Mta2* mutant, 440 genes had binding of all 3 factors (Mta2, Mbd3, and Zbtb7a) within the promoter region of the gene (Fig. 6C, D). GSEA analysis of these genes (Supp. Table 7) showed significant enrichment for oxidative phosphorylation (29 genes, p-value 4.44E-25, FDR q-value 2.22E-23) and adipogenesis (13 genes, p-value 1.22E-07, FDR q-value 3.05E-06), indicating that Zbtb7a and NuRD interact to regulate expression of genes in these pathways. Using an independent bioinformatics tool, we integrated our RNA-seq and ChIP-seq data using Binding and Expression Target Analysis (BETA) and identified 217 common down regulated target genes bound by all 3 factors (Mta2/Mbd3/Zbtb7a) [Supp. Table 8]. GSEA analysis of these common 217 genes revealed enrichment of oxidative phosphorylation (FDR q-value 3.56E-5), adipogenesis (FDR q-value 1.56E-3), and fatty acid metabolism (FDR q-value 1.69E-3) [Supp. Table 8], further validating our results that these pathways are directly regulated by NuRD/Zbtb7a.

### Human FSGS biopsies and Mta2 mutant kidneys exhibit common disease pathways

To explore the relationship of the findings in our *Mta2* mouse model to human FSGS, we examined changes in protein expression in glomeruli (G) and tubulointerstitium (TI) measured using spatial proteomics (Supp. Fig. 9A-D, Supp. Table 9). MTA2 protein was highly expressed in G and TI of human kidneys whereas the related homologues are weakly expressed (MTA1) or not detected (MTA3) in normal human kidney, indicating that MTA2 is the preferred NuRD component in the human kidney; all other NuRD components were also detected. We performed GSEA for proteins that showed a statistically significant change in expression in glomeruli in primary and secondary FSGS compared with reference nephrectomy tissues. This analysis revealed that, similar to RNA-seq results of *Mta2* mutant kidneys at 2 months of age, in human FSGS there was enrichment for downregulated proteins that function in OxPhos (FDR q-values 5.96E-10 for primary, 3.46E-8 for secondary FSGS) and fatty acid metabolism (FDR q-values 2.62E-8 for primary, 2.23E-4 for secondary FSGS) [Fig. 7A, B, Supp. Table 10]. GSEA of downregulated proteins in the TI of biopsies from primary and secondary FSGS also revealed enrichment for OxPhos (FDR q-values for primary 5.86E-7, secondary 7.91E-26) and fatty acid metabolism (FDR q-values for primary 8.54E-4, secondary 5.47E-8) [Fig. 7C, D, Supp. Table 10]. The complement and coagulation cascades were also enriched in the differentially expressed proteins in TI and glomeruli, with the most significant enrichment in the upregulated proteins of the secondary FSGS glomeruli (coagulation FDR q-value 3.16E-29, complement FDR q-value 1.57E-13) [Fig. 7A-C, Supp. Table 10]. Mitochondrion was the most significant Gene Ontology term for both primary FSGS (FDR q-value 3.92E-37) and secondary FSGS (FDR q-value 9.65E-52) in TI samples, (Supp. Table 11). Although there were far less upregulated genes than downregulated genes in our 2 month mutant RNA-seq data set, 77/217 genes that were upregulated in *Mta2* mutants at 2 months (>1.5fold, p< 0.05), had Mta2/Mbd3/Zbtb7a binding at the promoter (Fig. 6C). GSEA analysis of these 77 genes identified the complement (p-value 5.41E-03, FDR q-value 3.38E-02) and coagulation (p-value 1.91E-03, FDR q-value 3.18E-02) pathways as some of the most significant pathways (Supp. Table 12). RNA-seq and ChIP-seq integration using BETA for upregulated target genes revealed 115 common genes bound by Mta2/Mbd3/Zbtb7a. GSEA analysis showed enrichment in the complement (FDR q-value 2.52E-3) and coagulation (FDR q-value 4.22E-2) pathways, as well as allograft rejection (FDR q-value 2.52E-3) and epithelial to mesenchymal transition (FDR q-value 2.52E-3) [Supp. Table 13]. Overall, these results indicate that the *Mta2* mutant model of FSGS shares common disease pathways with human FSGS.

**Figure 7.**
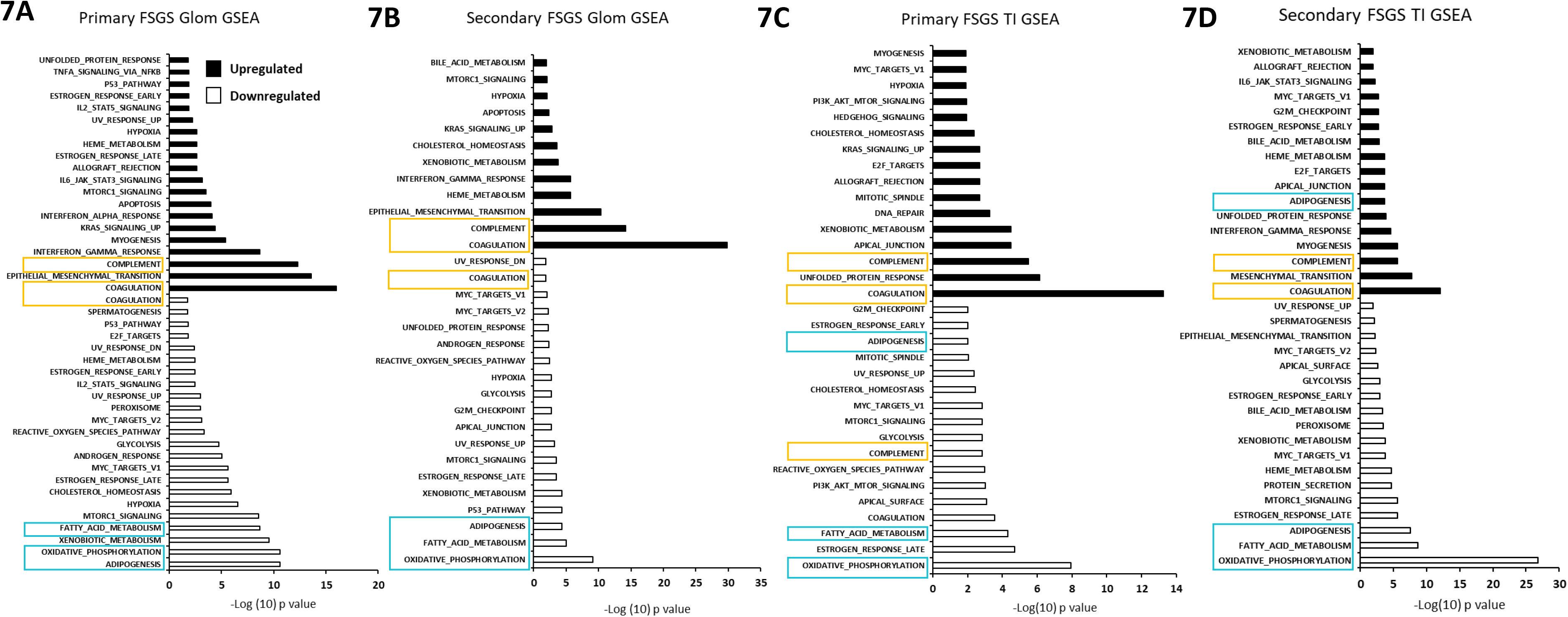
Human FSGS biopsies and *Mta2* mutant kidneys exhibit common disease pathways. **A-D.** Gene set enrichment analysis (GSEA) of differentially expressed proteins (p<0.05) in primary and secondary FSGS glomeruli (Glom) and tubulointerstitium (TI). Common pathways are present in downregulated genes in human FSGS and the *Mta2* mutant mouse (oxidative phosphorylation, adipogenesis, fatty acid metabolism), as well as upregulated genes (complement and coagulation).

## DISCUSSION

Epidemiological studies have shown than low nephron endowment is a significant risk factor for reduced GFR, albuminuria and CKD, with histological lesions of focal segmental glomerulosclerosis, tubulointerstitial fibrosis and inflammation (15, 20–23). However, the genetic and environmental factors that determine whether an individual with reduced nephron endowment at birth will develop CKD and whether disease will occur in childhood or in adult life are not well understood. A limitation in the field has been a lack of suitable animal models to study the molecular mechanisms of congenitally low nephron number.

We generated a mouse model with a moderate (∼30%) reduction in nephron number due to deletion of a core NuRD chromatin remodeling complex component, *Mta2*, in Six2+ nephron progenitor cells. In the developing embryo, maintenance and expansion of nephron progenitor cells and UB branching was not impaired in *Mta2* mutant metanaphroi. While induction of renal vesicles, the first epithelial tubule structure to form, was also not affected, their subsequent differentiation into mature nephron structures, such as glomeruli and proximal tubules, was reduced. The nephrons that formed in mutants are histologically indistinguishable from control kidneys, albeit in reduced numbers. However, transcriptional profiling at E17.5 revealed significant differences in gene expression between control and mutant mice. Comparison of single cell RNA-seq data from E18 embryonic kidneys revealed that while mutant nephrons had normal appearance by light microscopy, there was widespread reduction in expression in segment specific terminal differentiation genes that are required for the normal physiological functions of the kidney. In addition to evidence of impaired terminal differentiation, Gene Ontology (GO) analysis was highly significant for metabolic pathways among downregulated genes, including mitochondrial processes such as oxidative phosphorylation and lipid metabolism. While many studies have focused on the quantitative defect in nephrons as a risk factor for CKD, less attention has been given to the idea that the genetic or environmental factors that reduce nephron number may also impair their proper differentiation. We hypothesized that the failure of nephrons to fully achieve terminal differentiation and metabolic capabilities at the molecular level might persist in the post-natal period, which render the animal susceptible to injury and reduced renal function in the post-natal period.

*Mta2* mutant mice developed albuminuria at 4-6 weeks of age followed by histological evidence of FSGS at 8 weeks. There was progressive glomerulosclerosis and tubulointerstitial fibrosis, leading to reduced survival after 6 months of age. Glomerular enlargement was noted at 4 weeks of age (Supp. Fig. 7B-D), consistent with a recent report showing that mice with congenital low nephron number acquire glomerular hypertrophy between 2-6 weeks of age (4). Our model, and that of Good et al, share important phenotypic features, in particular, both had evidence of CKD in adolescence and early adulthood. Our findings are also consistent with those seen in children who have a history of preterm birth or very low birth weight who were found to have albuminuria, glomerular hypertrophy and glomerulosclerosis on renal biopsy (24).

To investigate the mechanism of FSGS in our model of congenital low nephron number, we performed RNA-seq at 2 months of age, when FSGS was first apparent by renal histopathology. We found reduced expression of segment specific terminal differentiation genes, similar to transcriptional profiling of embryonic kidneys. GO analysis also revealed that downregulated genes were highly enriched for genes required for mitochondrial metabolic functions, with oxidative phosphorylation and fatty acid metabolism being very prominent pathways among altered genes. Therefore, the disrupted pattern of gene expression established in late embryogenesis persisted in the post-natal kidney. Based on these results, we tested for defects in lipid metabolism and mitochondrial function. Our studies revealed neutral lipid accumulation in glomeruli as early as 4 weeks of age, prior to development of FSGS lesions, whereas impaired mitochondrial dysfunction was not detected in the renal cortex until 3 months of age. These results suggest a model in which impaired fatty acid oxidation leads to glomerular injury and subsequent development of FSGS. In this paradigm, mitochondrial dysfunction is the result, not the cause of impaired fatty acid metabolism, and contributes to the progression of kidney injury and fibrosis observed in the *Mta2* mutants after 2 months of age. Our results are consistent with prior studies in which impaired fatty acid oxidation, lipid accumulation and mitochondrial dysfunction have been implicated in the pathogenesis of CKD progression, and glomerular disease in humans and animal models (25–27). Our regional proteomics data from human kidney biopsy samples provide direct experimental evidence that disrupted mitochondrial metabolism, affecting both glomeruli and the tubulointerstitium, plays a role in FSGS. In addition, similar to the data in our mouse model, complement and coagulation pathways were enriched in human FSGS biopsies. While these pathways have not been typically linked to FSGS, our studies suggest they may warrant further investigation in this disease.

A limitation of our studies is the possibile inability to detect earlier changes in mitochondrial function restricted to specific cellular compartments, such as podocytes, because we performed respirometry on renal cortical slices. However, whether impaired fatty acid oxidation or mitochondrial dysfunction is the initial inciting event, these results strongly suggest that the metabolic demands of post-natal somatic growth and maturation represent a susceptible period for the kidney, when congenital low nephron number is present. In support of this is idea, marked glomerular enlargement was associated with increased body surface area in males with low nephron number, thereby linking metabolic demand to adaptive hypertrophy (28). In addition, metabolic stress, in the form of postnatal overfeeding or high fat diet superimposed on low nephron endowment resulted in albuminuria and glomerulosclerosis in animal models (29–35). The mouse kidney undergoes rapid growth between 2-6 weeks of age and reaches near adult size by 8 weeks (36). During this period of rapid growth, the kidney relies heavily on fatty acid as a source of energy (36, 37). We propose that in the face of impaired fatty acid oxidation, the *Mta2* mutant kidney cannot meet the metabolic demands of post-natal somatic growth.

Consequently, the compensatory hypertrophy becomes maladaptive leading to development of CKD at 4-8 weeks of age. A recent study showed that mitochondrial dysfunction plays a critical role in developmental programming of cardiovascular disease and HTN (38). While a role for mitochondrial dysfunction has also been suggested for fetal programming of CKD, it has not been demonstrated experimentally (39). The findings in the *Mta2* mutant model of congenital low nephron number support a role altered mitochondrial fatty acid metabolism in developmental programming of kidney disease.

One of the major mechanisms by which cells respond to stress or injury is by utilizing the chromatin modifying machinery to reprogram gene regulatory networks. The Nucleosome Remodeling and Deacetylase (NuRD) chromatin-remodeling complex fine-tunes the activity of regulatory elements, making them particularly responsive to changes in the state (healthy vs. injured) of the cell. We investigated whether genes with altered expression in the mutant were direct targets of NuRD. ChIP-seq studies revealed that Mta2 and another core NuRD component, Mbd3, bound the promoters of many differentially expressed genes, including a large number of metabolic genes that were downregulated. This observation is consistent with bioinformatics predictions from the MitoCarta data set that placed Mta2 in the neighborhood of transcriptional regulators of nuclear genes encoding mitochondrial proteins (17). Our study is the first to our knowledge that provides direct experimental evidence that Mta2-NuRD is likely a direct regulator of genes required for mitochondrial metabolism. Our bioinformatics analyses indicated that Zbtb7a/b, cooperated with NuRD to regulate metabolic genes, which was also supported by our ChIP-seq studies. Zbtb7a/b has not previously been implicated in kidney injury. Zbtb7a/7b are members of a larger family of factors that contain C-terminal C2H2/Kruppel-type zinc fingers that bind DNA and an N-terminal BTB (broad complex, tram track, bric-a-brac) domain that mediates interaction with transcriptional co-regulators [reviewed in (40)]. Zbtb7a and 7b have highly conserved Zn fingers in their C-termini and are thought to interact with very similar DNA motifs (41). Genetic studies in mice indicated that Zbtb7a and 7b have both redundant and unique functions in T cells [reviewed in (40)]. To validate our results in human tissues, we analyzed KPMP data for transcription factor binding site enrichment for genes that showed differential expression and chromatin accessibility in proximal tubules in the adaptive state (aPT). The aPT was associated with progression of CKD and acute kidney injury (AKI) to CKD transition (42). This analysis revealed that ZBTB7A DNA motifs were significantly enriched in aPT target genes (adjusted p 2.74 X 10E-6) (42). Single cell RNA-seq data from human CKD samples obtained from a different cohort of patients revealed that ZBTB7A/B expression was increased in podocytes in human CKD samples (43). Together, these independent human data sets provide validation for a role for ZBTB7A/B in human CKD, as suggested by analysis of our mouse model.

Because epigenetic changes mediated by chromatin remodeling complexes, such as NuRD, and their associated transcription factors are sensitive to environmental insults, they are expected to play an important role in developmental programming. Importantly, the reversible nature of these chromatin modifications makes them potentially attractive targets for therapeutic interventions to reprogram gene expression to prevent the transition from compensatory hypertrophy to CKD. Drugs targeting enzymes that modify chromatin have already entered the clinic for the treatment of several cancers and could potentially be used for preventing the maladaptive consequences of glomerular hypertrophy that promote the transition to CKD.

## METHODS

### Animals

All animal studies were carried out in accordance with the guidance of the Institutional Animal Use and Care Committee of Washington University School of Medicine. *Mta2* homozygous floxed (*Mta2^flx/flx^*) mice were a generous gift from Dr. Yi Zang, University of North Carolina Chapel Hill and were generated as described (44). *Mta2^flx/flx^* mice were bred with *Six2*-TGC BAC transgenic mice that express Cre recombinase in the Six2+ nephron progenitor cells (45) to generate *Six2Cre Mta2flx/flx* mutants. For genotyping, genomic DNA was amplified with the following oligonucleotide primers: 5′-GCTGAAGCAGACAGCAAAC-3′ and 5′- CATGCCAGGTTTTGAACCC-3’.

### Tissue Preparation, Histology, Electron Microscopy and Immunofluorescence

Kidneys were harvested and divided on the transverse plane into portions for flash freezing (RNA/protein) or fixing with 10% buffered formalin (histological staining) and 4% PFA (antibody staining and Oil Red O staining). Paraffin sections (4 µm) were stained with hematoxylin/eosin, periodic acid Schiff (PAS) or Masson’s trichrome (Saint Louis University Research Microscopy and Histology Core) and imaged with bright field illumination. For antibody staining and Oil Red O staining (Saint Louis University Research Microscopy and Histology Core), PFA fixed kidney portions were mounted in OCT and frozen in a 2-methylbutane/dry ice and ethanol bath. 7 μm frozen sections were stained with primary antibodies and conditions (Supp. Table 14). Reactivity was detected using fluorescently labeled secondary antibodies (Supp. Table 14). Sections were counterstained with DAPI (Sigma Aldrich), mounted in Mowiol 4-88 (Poly Sciences), and digital images acquired using a Nikon 80i epifluorescence microscope and Retiga R3 camera. For electron microscopy, small pieces ∼1-3 mm^3^ of kidney cortex from 8 week old mice were fixed in 2.5% glutaraldehyde fixative (2.5% glutaraldehyde, 0.1M sodium cacodulate pH 7.2-7.25, 2% sucrose, 2mM CaCl2) and processed by Saint Louis University Research Microscopy and Histology Core. Images were acquired using a JEOL 1400 Plus transmission electron microscope.

### Semi-quantitative Tubulointerstitial damage and glomerulosclerosis

Average semi-quantitative tubulointerstitial damage was evaluated by determining the % damage per high powered field (40X) (Grade 0, no change; Grade 1, <25%; Grade 2, 25-50%; Grade 3, 50-75%; Grade 4, >75%). At least 16 individual PAS stained images were evaluated from 3 biological replicates for each genotype. Average semi-quantification of glomerulosclerosis was evaluated by determining the % sclerosis per glomeruli (Grade 0, no change; Grade 1, <25%; Grade 2, 25-50%; Grade 3, 50-75%; Grade 4, >75%). At least 40 PAS stained glomeruli were evaluated from 3 biological replicates for each genotype. Statistical analysis was performed using a two-tailed Student’s t-test. H&E, PAS, and Masson’s Trichrome stained kidney sections from 4 week and 8 week control and mutant mice were examined blinded by two independent pathologists who both independently diagnosed the mutant mice with FSGS.

### Urinary Albumin

Spot urine collections were stored frozen at -80 until analysis, then thawed on ice and debris pelleted. 10ul was used to estimate presence of protein (Siemens Multistix 5, product no. 2309) before analysis. Albumin was measured using Ethos Biosciences Exocell Albuwell M murine microalbuminuria kit (product no. 1011) according to the manufacturer’s protocol. Absorbance was read at 450nm on a Chromate 4300 plate reader. Urine samples were diluted to fit the standard curve. Albumins were normalized to creatinine determined by the Creatinine Companion (Ethos Biosciences Exocell, product no. 1012), absorbance read at 500nm. A minimum of 2 replicates per mouse were averaged. Results are reported as the Log of the Albumin Creatinine Ratio (ACR). Statistical analysis was performed using multiple paired T tests, Holm-Sidak method (Graphpad Prism software).

### Quantitation of ureteric bud branching, cap mesenchyme and differentiating nephron structures

Embryonic day 17.5 kidneys were immunostained for Six2, Cytokeratin and DAPI. Six2+ caps and Cytokeratin positive UB tips were counted from at least 12 non-sequential sections (10× magnification) from at least 3 independent embryos for each genotype. Results were reported as the average number of caps or UB tips per high-powered field (HPF) ± SEM. For renal vesicles, embryonic day 17.5 kidneys were immunostained for NCAM and Lef1. For terminal nephron segments, immunostaining was performed to detect Wt1 (glomeruli), podocin (glomeruli), megalin (proximal tubule), NKCC2 (loop of Henle), NCC (distal tubule) and cytokeratin (collecting duct). All structure quantification was performed by counting positive staining for cross sections of tubules, e.g. for proximal tubules multiple cross-sections of the same tubule may have been counted, however counting was performed consistently for both control and mutant samples. Structures were counted on at least 12 non-sequential sections (10X magnification) from at least 3 independent embryos for each genotype. Statistical analysis was performed using a two-tailed Student’s t-test. Glomeruli were counted using the acid maceration method (2) in at least 2 independent biological replicates for each genotype. Statistical analysis was performed using a one way Anova with Tukey’s multiple comparison test.

### Cell Proliferation and Apoptosis

Mitotic index of nephron progenitor cells was determined by staining embryonic kidneys at 17.5 for pHH3 and Six2. Nuclei were stained using DAPI. The total number of pHH3 and Six2/pHH3+ cells were counted on at least 36 images from 12 non-sequential sections (20× magnification) from at least three independent embryos for each genotype. FACS isolation of Six2-GFP+ cells and cell cycle profiles were determined as previously described (46). Apoptosis was determined by performing TUNEL analysis using the ApopTag Red *In Situ* Apoptosis Detection Kit (Millipore). The total number of TUNEL positive cells were quantified per high powered field (HPF) in addition to the total number of Six2/TUNEL, Wt1/TUNEL, and LTL/TUNEL+ cells/HPF. For all stages at least 24 HPF for each genotype was quantified from at least 12 non-sequential sections from at least 3 biological replicates for each genotype. Results are reported as the average # of cells/HPF ± SEM. Statistical analysis was performed using a two-tailed Student’s t-test.

### RNA-sequencing (RNA-seq)

Total RNA was isolated from kidneys of three independent biological samples for each age (E17.5, 2 months) and genotype using an RNeasy Mini Kit (Qiagen 74104) with on-column DNAse I treatment. Ribosomal RNA depletion, barcoded library construction and sequencing were performed as previously described (47). Reads were aligned to the mouse mm10 genome using the TMAP aligner map4 algorithm. Soft-clipping at both 5′ and 3′ ends of the reads was permitted during alignment to accommodate spliced reads, with a minimum seed length of 20 nt. Genome-wide strand-specific nucleotide coverages were calculated from the aligned bam files for each sample using the *genomecoveragebed* program in BEDTools (48) and the nucleotide coverage for all non-redundant exons for each gene were summed using custom R scripts (49). Normalization factors were calculated by averaging the total exon coverage for all replicates and dividing this average by the total exon coverage for each individual sample. The total coverage for each gene in each replicate was then multiplied by these factors after adding an offset of 1 to each gene to preclude division by 0 in subsequent calculations. The averages and p values of the coverage values for all genes in the individual groups were calculated using Microsoft Office Excel. The expression values for each gene are the normalized strand-specific total nucleotide coverage for each gene. Volcano plots created using EnhancedVolcano in R.

### Chromatin Immunoprecipitation Sequencing (ChIP-seq)

ChIP-seq was performed as previously described (47). Two month old kidneys were harvested and isolated kidney cortex was cross-linked with 2 mM Di (N succinimidyl) glutarate (DSG, Proteo Chem C1104) in PBS for 30 min followed incubation in 1% formaldehyde (Thermo Scientific 28906) for an additional 10 min at room temperature and quenched with 125 mM glycine for 5 min. Pre-cleared chromatin was incubated with primary antibody (Supp. Table 14) overnight with rocking at 4 °C. Complexes were isolated using protein A/G magnetic beads and eluted DNA was isolated for library construction. Libraries were made using NEBNext Ultra II Library Kit (E7645) and sequenced on an Illumina Nova Seq at a depth of 100 million paired end reads. Reads were aligned to the mm10 mouse genome using the Bowtie2 read aligner.

Genome coverage was calculated for ChIP and input samples using Deeptools bamCoverage. ChIP peaks were called using Macs2 standard peak calling parameters using input samples as control. Called peaks were assigned to genomic features (promoters, enhancers, introns, or exons) using Bedtools based on peak overlap with a defined feature. Genomic features (active/inactive promoters and predicted enhancers) were defined using published ENCODE (50) datasets for H3K4me3 (ENCSR000CAN), H3K27ac (ENCSR000CDG), H3K4me1 (ENCSR000CAF) and DNase-seq (ENCSR000CNG) as previously described (47). ChIP-seq was performed on two biological replicates for each genotype and antibody tested. Gene Ontology analysis was performed using the ChIPseeker R package, gProfiler, and Gene Set Enrichment Analysis (51, 52). Binding sites were compared to RNA-seq data sets using Binding and Expression Target Analysis, BETA (53). Venn diagrams were created using Venny (54).

ChIP-seq tracks were made using IGV.

### High resolution respirometry (OROBOROS)

Oroboros was carried out according to (55). Briefly, kidneys were harvested and immediately placed in ice cold BIOPS buffer (10 mM EGTA, 50 mM MES, 0.5 mM DTT, 6.56 mM MgCl2, 5.77 mM ATP, 20 mM imidazole, and 15 mM phosphocreatine, pH 7.1) until tissue preparation. Prior to permeabilization, 1mg pieces of kidney cortex were isolated and blotted dry (3 times), weighed (1mg approximately) and placed in the OROBOROS corresponding chambers containing mitochondrial respiration solution (MiR05, 0.5 mM EGTA, 3 mM MgCl2, 60 mM k-lactobionate, 20 mM taurin, 10 mM KH2PO4, 20 mM HEPES, 110 mM sucrose, and 1 g/l BSA, pH 7.1). Digitonin (2 µM) was added to the solution to permeabilize the tissue. Oxygen was injected into each chamber maintaining the oxygen levels around 400 pmol during the assay. The following substrates were injected sequentially into the chamber: 1.5 mM octanoyl-L-Carnitine (OC), 5mM Pyruvate and 2mM Malate (Pyr/Mal), 10mM Glutamate (G), 20 mM adenosine diphosphate (ADP), 20 mM succinate (Succ), and 0.5 µM FCCP (3 pulses). A period of stabilization followed the addition of each substrate, and the oxygen flux per mass was recorded using DatLab 6.1 software (OROBOROS Instruments). Results are reported as the average oxygen flux per mass ± SEM. For 2 and 3 month experiments at least 10 biological replicate experiments were performed for each genotype. Statistical analysis was performed using multiple paired t-tests.

### Quantification of glomerular perimeter and area

4 week old mouse kidney cortex was isolated and glomeruli purified using the differential adhesion method (56). Briefly, the cortex is minced and disaggregated with collagenase V solution (1 mg/mL) for 15 minutes in a 37°C water bath. The digestion is stopped with 10% FBS in DMEM then washed with HBSS and filtered through a 150 mesh strainer, and filtrate then passed through a 200 mesh strainer. Filtrate from the 200 mesh strainer is passed through a 40 µm cell strainer and the obtained fragments from the 40 µm strainer is obtained by inverting the strainer and flushing with HBSS. Fragments in HBSS from the 40 µm strainer is placed in a 10 cm plate (430167, Corning) and tissue allowed to settle for 2 minutes. Tubules are adherent to the plate and glomeruli float. Glomeruli are pulled off of the plate using a large disposable transfer pipette and put through a second 40 µm strainer. Fragments are obtained from the strainer again by inverting and flushing with HBSS. Obtained fragments are placed on a second 10 cm plate and allowed to sit for 2 minutes. Glomeruli in HBSS are pulled off of the plate and added to a 50 mL conical with 0.01% (m/v) PVP and pelleted for 5 minutes at 600 g at 4°C.

Glomeruli were re-suspended in PBS +Ca, +Mg with 1% FBS, plated on a cell culture plate, and imaged on a Leica DMi1 microscope with a Leica MC170 HD 5 MP HD camera. Glomerular area was measured using Image J by first calibrating pixel size with a scale bar. The perimeter of each glomeruli was measured using the freehand selection tool, and the area of the ROI was calculated and reported as µm^2^. 111 glomeruli were analyzed from 2 biological replicates for *Mta2flx/flx* and 197 glomeruli from 4 *Six2Cre Mta2flx/flx* biological replicates. Statistical analysis was performed using an unpaired Student’s t-test.

### Proteomic analysis of human nephrectomy and kidney biopsy tissue

#### Human subjects study approval and patient characteristics

Formalin-fixed paraffin embedded (FFPE) kidney biopsy samples were obtained from transplant donor kidneys (n=4), primary FSGS (n=5), and secondary FSGS (n=7). The institutional review board at Ohio State University approved human subject studies performed.

#### Sample preparation for mass spectrometry

The tissue collection and proteomics studies were done in two batches. Batch 1 included transplant donor kidneys (n=4), minimal change disease (n=9), primary FSGS (n=3) and secondary FSGS (n=4) and batch 2 included primary FSGS (n=2) and secondary FSGS (n=3). For laser capture microdissection (LCM), 10 μm sections were obtained from paraffin blocks onto PEN membrane slides (Zeiss**). Sections were deparaffinized with xylene and rehydrated in ethanol gradient to mass spectrometry grade water. Sections were then treated with 70%, then 90% ethanol, followed by two changes in 100% ethanol for 2 minutes for each step. The glomerular and tubulointerstitial compartment was collected by using LCM system (PALM Technologies, Carl Zeiss Micro Imaging GmbH, Munchen, Germany) containing a PALM MicroBeam and RoboStage for high throughput sample collection and a PALM RoboMover (PALM Robo software, version 2.2, Carl Zeiss, Microimaging GmbH, Munchen, Germany).

Tissue sections were captured into 0.5% Rapigest (Waters Corporation, Milford, MA) in 50 mM sodium bicarbonate and stored at -80°C until protein retrieval. For protein extraction, samples were boiled for 20 minutes and incubated at 60°C water bath for 2 hours. For protein digestion, trypsin (Promega, Madison, WI) was added at ratio of 1:30 (trypsin:protein) assuming that 2 μg protein was recovered from 10,000 cells. Protein-trypsin mix was incubated overnight at 37°C followed by incubation with formic acid at final concentration of 30-40% for 30 minutes minute at room temperature to completely precipitate the Rapigest. Samples were centrifuged at 15,000 rpm at 4°C for 10 minutes to remove the cell debris and dried using speedvac. Dried peptides were solubilized by resuspending in 20 μl of 2% acetonitrile with 0.1% formic acid followed by sonication for 1-2 minutes in 4°C water bath for 10 minutes. Final peptide concentration was measured by absorbance at 280 nm on a Nanodrop ND-2000 spectrometer (Thermo Fisher, Waltham, MA).

#### Proteomics by liquid chromatography and mass spectrometry

Liquid chromatography tandem-mass spectrometry (LC-MS/MS) analysis was performed with a Thermo Scientific Easy1200 nLC (Thermo Scientific, Waltham, MA) coupled to a tribrid Orbitrap Eclipse (Thermo Scientific, Waltham, MA) mass spectrometer. In-line de-salting was accomplished using a reversed-phase trap column (100 μm × 20 mm) packed with Magic C18AQ (5-μm 200Å resin; Michrom Bioresources, Auburn, CA) followed by peptide separations on a reversed-phase column (75 μm × 270 mm) packed with ReproSil-Pur C18AQ (3-μm 120Å resin; Dr. Maisch, Baden-Würtemburg, Germany) directly mounted on the electrospray ion source. A 180-minute gradient using a two-mobile-phase system consisting of 0.1% formic acid in water (A) and 80% acetonitrile in 0.1% formic acid in water (B). The chromatographic separation was achieved over a 180 min gradient from 8 to 30% B over 180 min, 30 to 45% B for 10 minutes, 45 to 60% B for 3 minutes, 60 to 90% B for 2 mininutes and held at 90% B for 10 minutes at a flow rate of 300 nL/minute. A spray voltage of 2300 V was applied to the electrospray tip in line with a FAIMS source using varied compensation voltage -40, -60, -80 while the Orbitrap Eclipse instrument was operated in the data-dependent mode, MS survey scans were in the Orbitrap (Normalized AGC target value 300%, resolution 120,000, and max injection time 50 ms) with a 3 second cycle time and MS/MS spectra acquisition were detected in the linear ion trap (Normalized AGC target value of 50% and injection time 20 ms) using HCD activation with a normalized collision energy of 27%. Selected ions were dynamically excluded for 60 seconds after a repeat count of 1.

#### Mass spectrometry data processing

All raw MS data were processed using Proteome Discoverer (v2.5.0.400, Thermo Fisher Scientific). Spectra were searched against the human proteome obtained from Uniprot (August 2022)(57) using Sequest HT with a fixed modification of carbamidomethylation of cysteine and variable of modifications of oxidation of methionine, oxidation of proline and N-terminal Acetylation. Tryptic peptides with a maximum were allowed, with precursor and fragment ion tolerances set at 10ppm and 0.6 Da, respectively. The false discovery rate (FDR) for both peptide and protein identification were set at 0.01. Volcano plots created using EnhancedVolcano in R. Venn diagram created using BioTools.fr.

## ACKNOWLEDGEMENTS

We thank Noemi Arias for her technical assistance in support of Oroboros experiments. Dr. Parikh was supported by an OSU Department of Internal Medicine Junior Investigator Award (SVP). Dr. Basta was supported by an American Society of Nephrology Transition to Independence Grant. Dr. Rauchman was supported by the Washington University Chromalloy Renal Diseases Endowment, 5R01DK12987902, VA Merit Award BX-003674, and Pediatric Center of Excellence in Nephrology 5P50DK133943.

## MATERIALS & CORRESPONDENCE

Please address data and materials requests to Michael Rauchman.

## DATA AVAILABILITY

All data is in the process of being made publicly available in GEO.

## Notes

### Competing Interest Statement

The authors have declared no competing interest.

