## Supplemental Figures for "Deletion of NuRD component *Mta2* in nephron progenitor cells causes developmentally programmed FSGS"

# S1

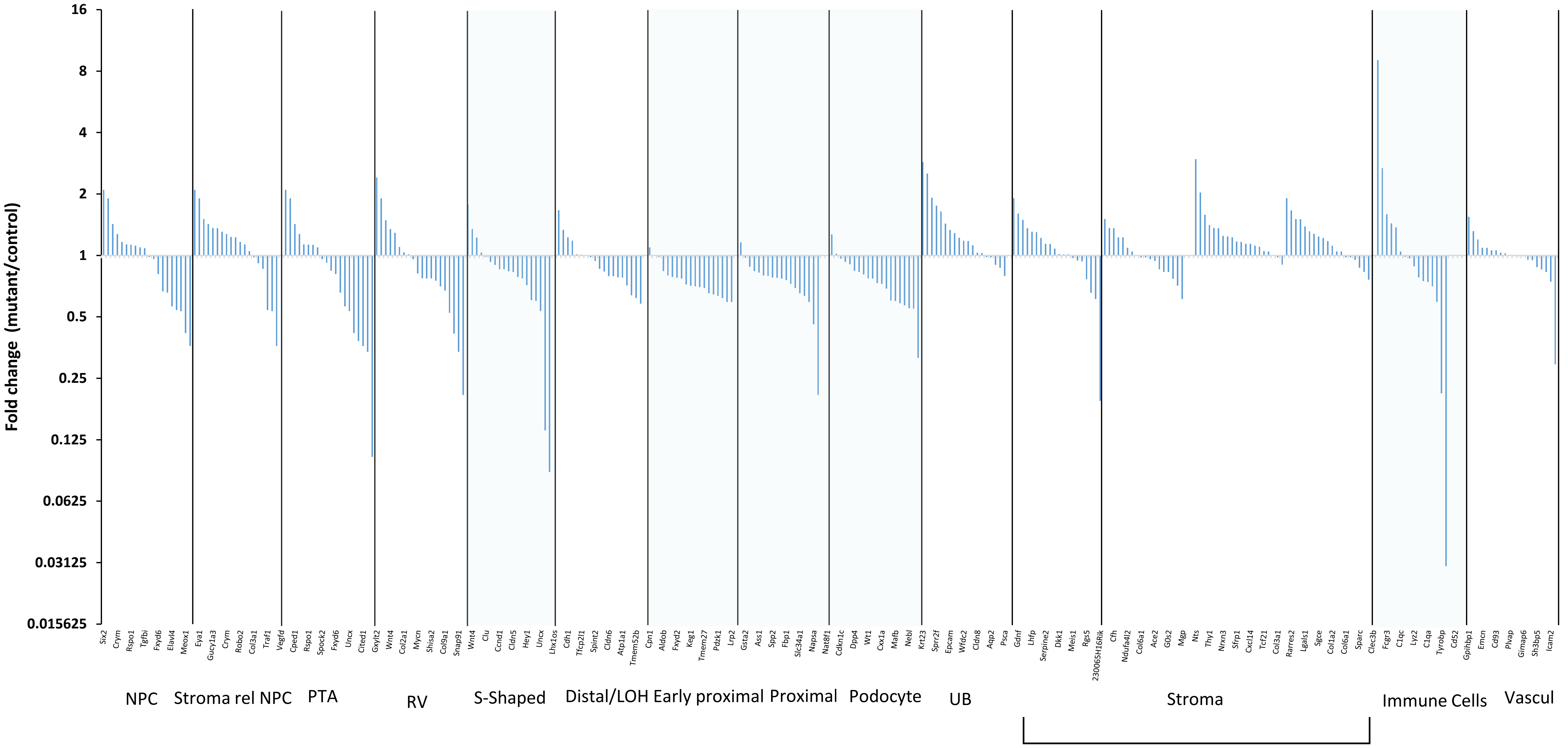
Fold change (mutant/control)
16
8
4
2
1
0.5
0.25
0.125
0.0625
0.03125
0.015625
Six2 Crym Rspo1 Tgfbi Fxyd6 Elavl4 Meox1
Eya1 Gucy1a3 Crym Robo2 Col3a1 Traf1
Vegfd Cped1 Rspo1 Spock2 Fxyd6 Uncx Cited1
Gxylt2 Wnt4 Col2a1 Mycn Shisa2 Col9a1 Snap91
Wnt4 Clu Ccnd1 Cldn5 Hey1 Uncx
Lhx1os Cdh1 Tfcp2l1 Spint2 Cldn6 Atp1a1 Tmem52b
Cpn1 Aldob Fxyd2 Keg1 Tmem27 Pdzk1 Lrp2
Gsta2 Ass1 Spp2 Fbp1 Slc34a1 Napsa
Nat8f1 Cdkn1c Dpp4 Wt1 Cxx1a Mafb Nebl
Krt23 Sprr2f Epcam Wfdc2 Cldn8 Aqp2 Psca
Gdnf Lhfp Serpine2 Dkk1 Meis1 Rgs5 230065H16Rik
Cfh Ndufa4l2 Col6a1 Ace2 G0s2 Mgp Nts Thy1 Nrxn3 Sfrp1 Cxcl14 Tcf21 Col3a1 Rarres2 Lgals1 Sgce Col1a2 Col6a1 Sparc
Clec3b Fcgr3 C1qc Lyz2 C1qa Tyrobp Cd52
Gpihbp1 Emcn Cd93 Plvap Gimap6 Sh3bp5 Icam2
NPC
Stroma rel NPC
PTA
RV
S-Shaped
Distal/LOH
Early proximal
Proximal
Podocyte
UB
Stroma
Immune Cells
Vascul

S2

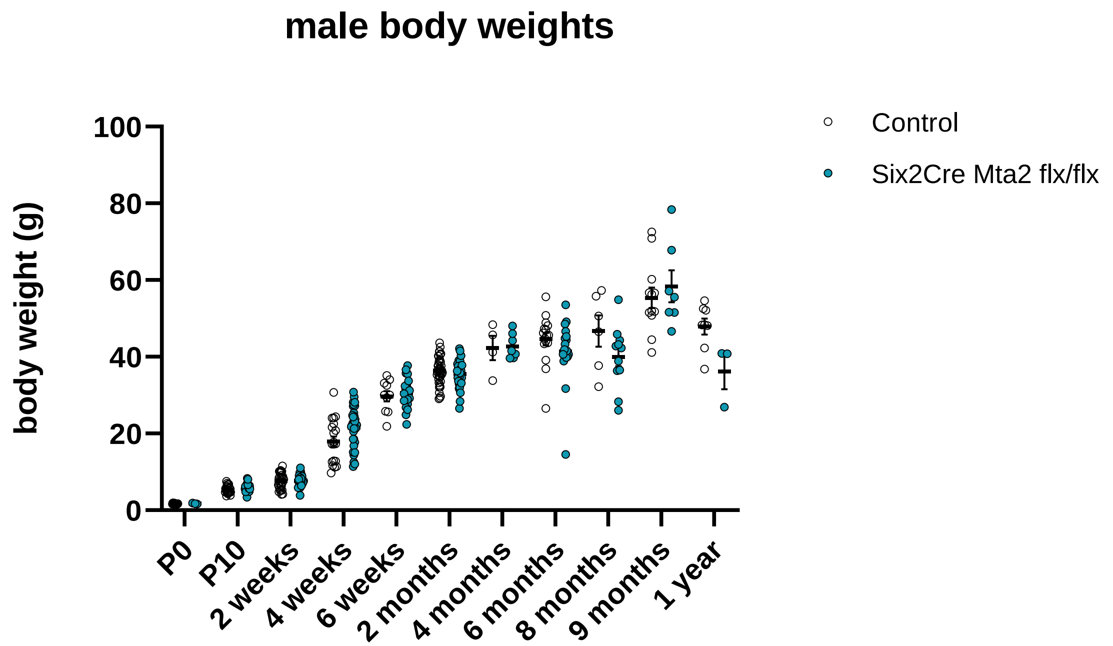
male body weights
body weight (g)
100
80
60
40
20
0
P0
P10
2 weeks
4 weeks
6 weeks
2 months
4 months
6 months
8 months
9 months
1 year
Control
Six2Cre Mta2 flx/flx

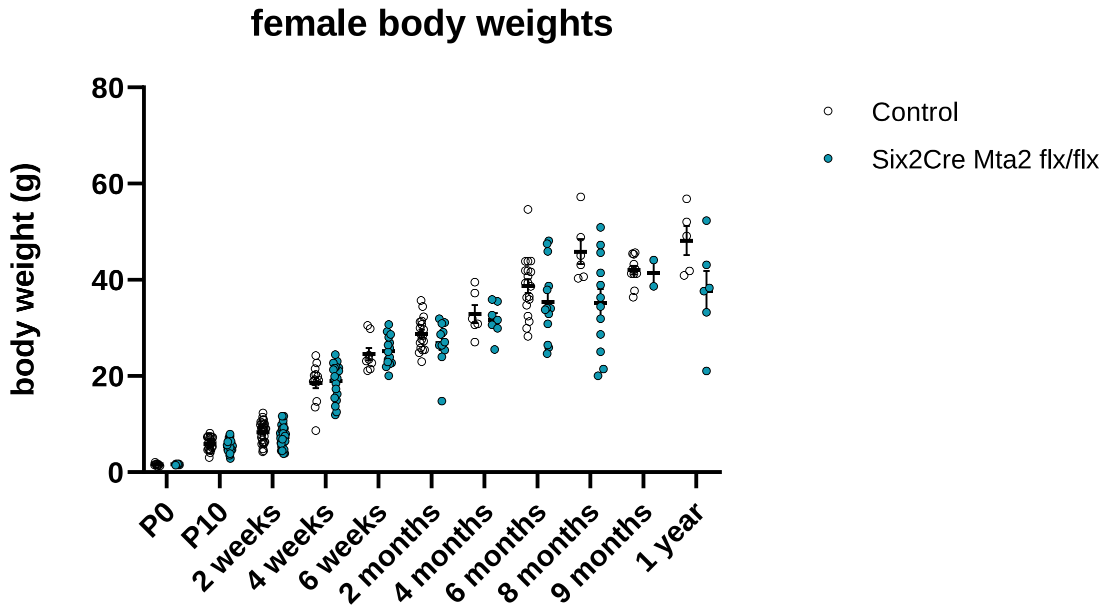
female body weights
body weight (g)
80
60
40
20
0
P0
P10
2 weeks
4 weeks
6 weeks
2 months
4 months
6 months
8 months
9 months
1 year
Control
Six2Cre Mta2 flx/flx

S3

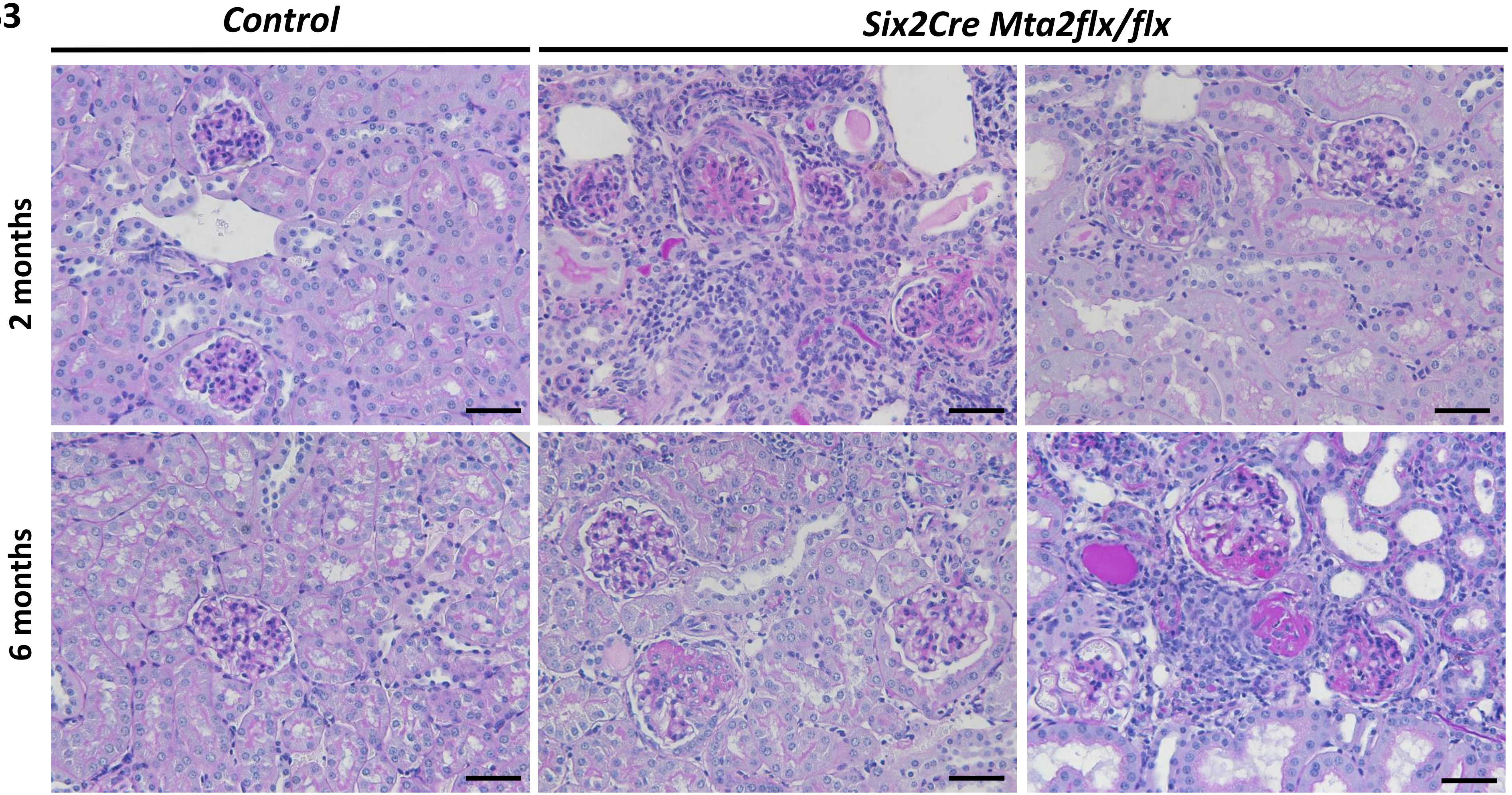

S4

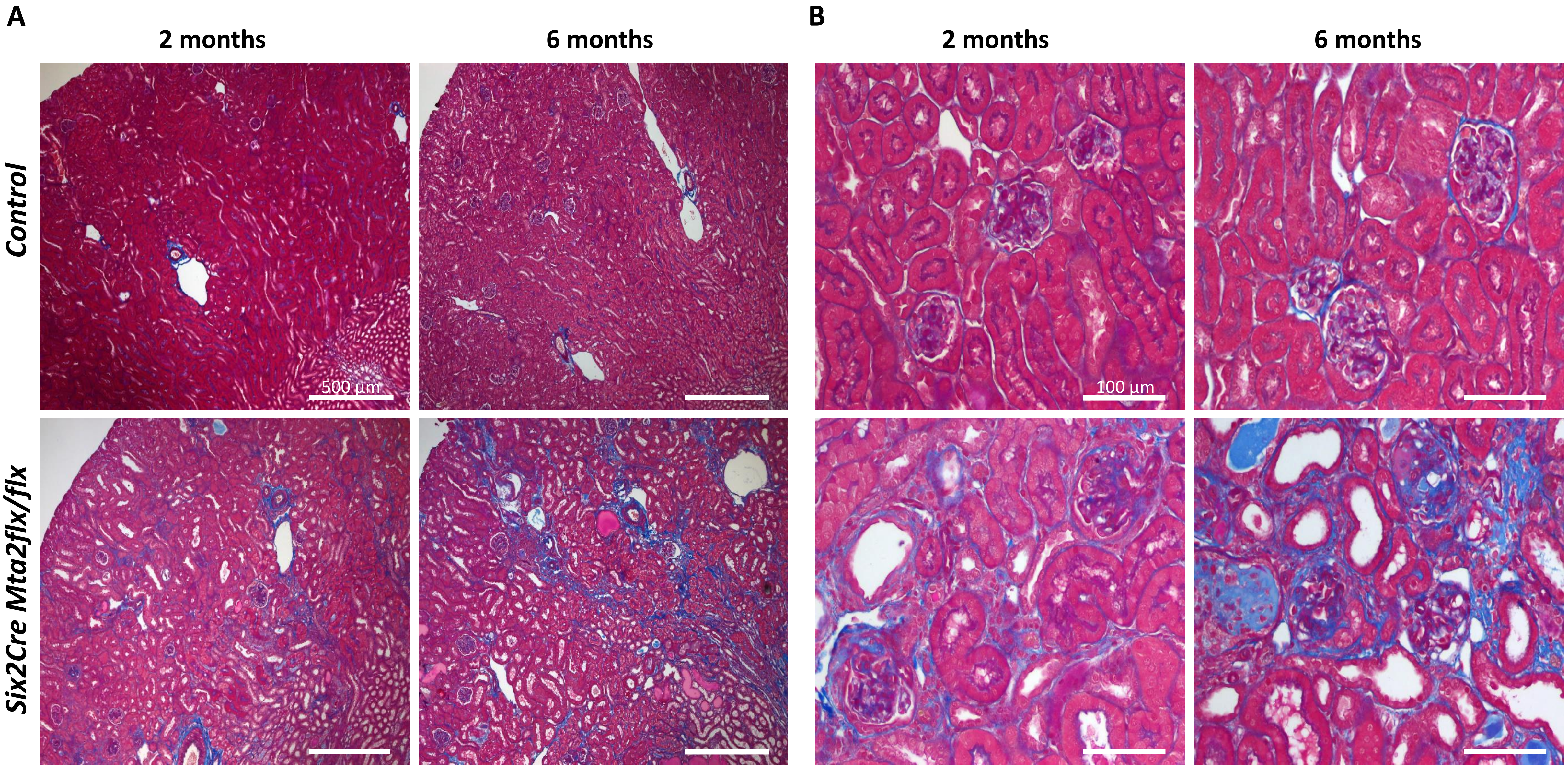
A
B
2 months
6 months
2 months
6 months
Control
Six2Cre Mta2flx/flx
500 µm
100 µm

S5

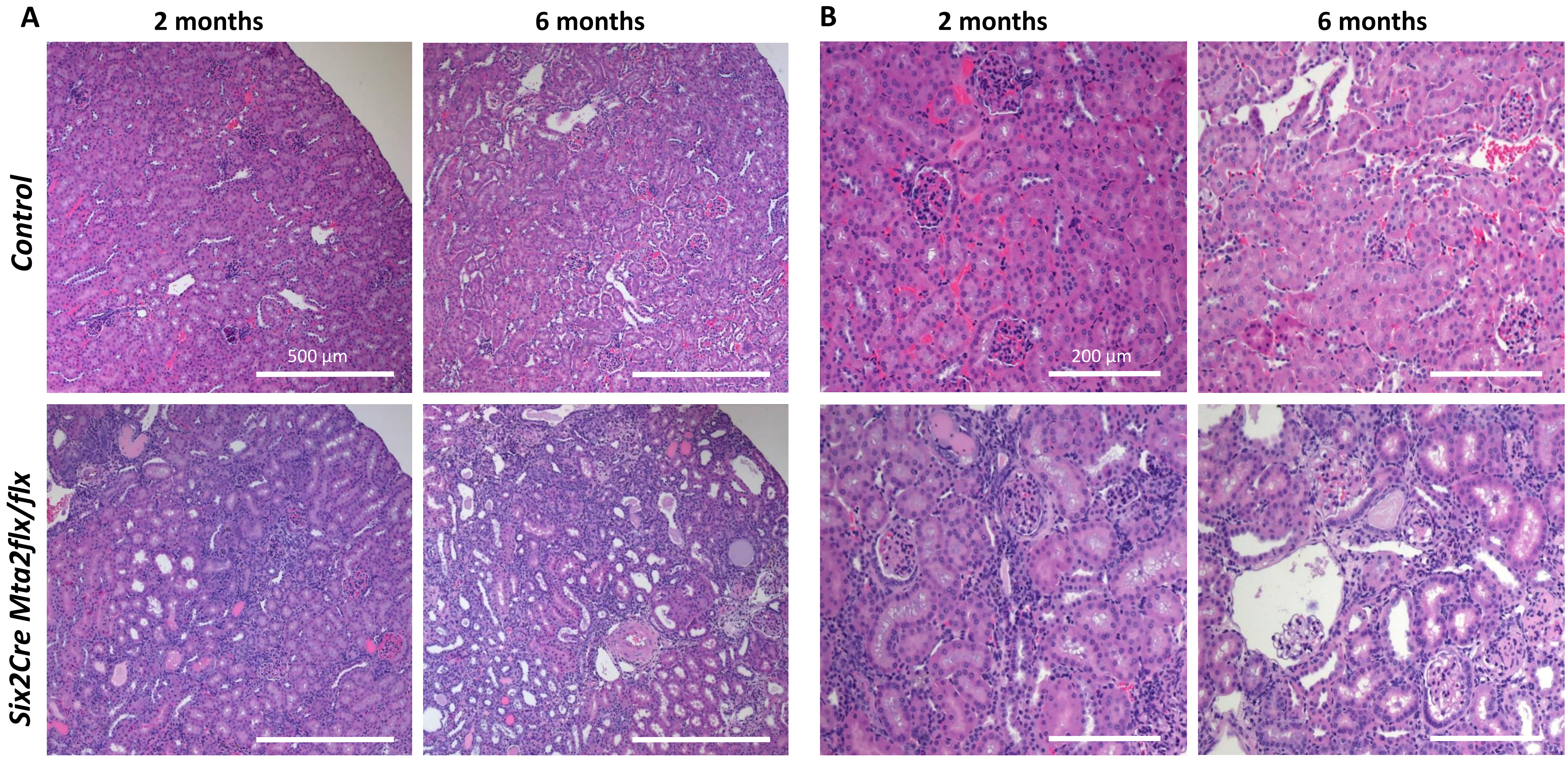
A
2 months
6 months
B
2 months
6 months
Control
Six2Cre Mta2flx/flx
500 µm
200 µm

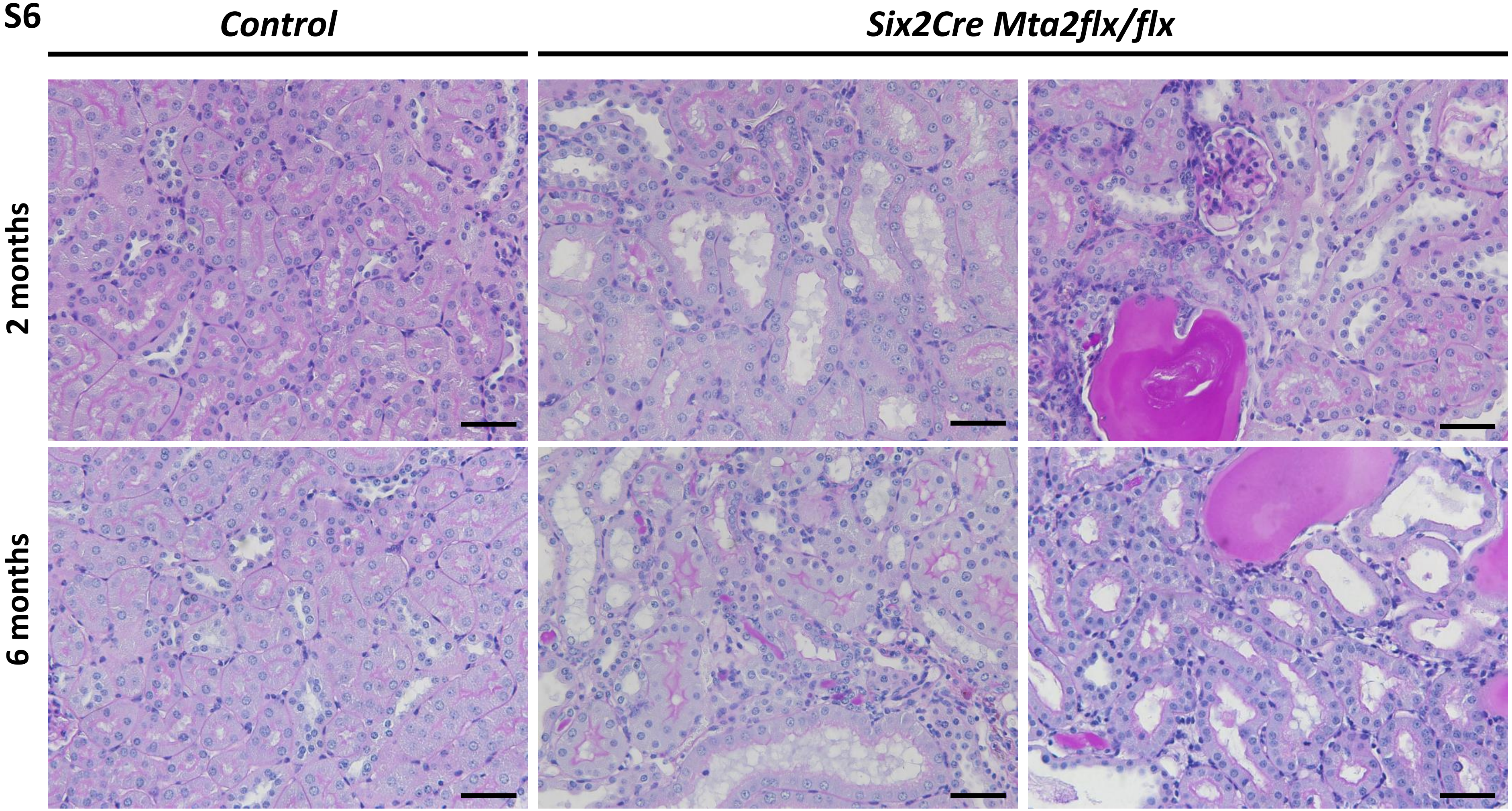
S6
Control
Six2Cre Mta2flx/flx
2 months
6 months

S7

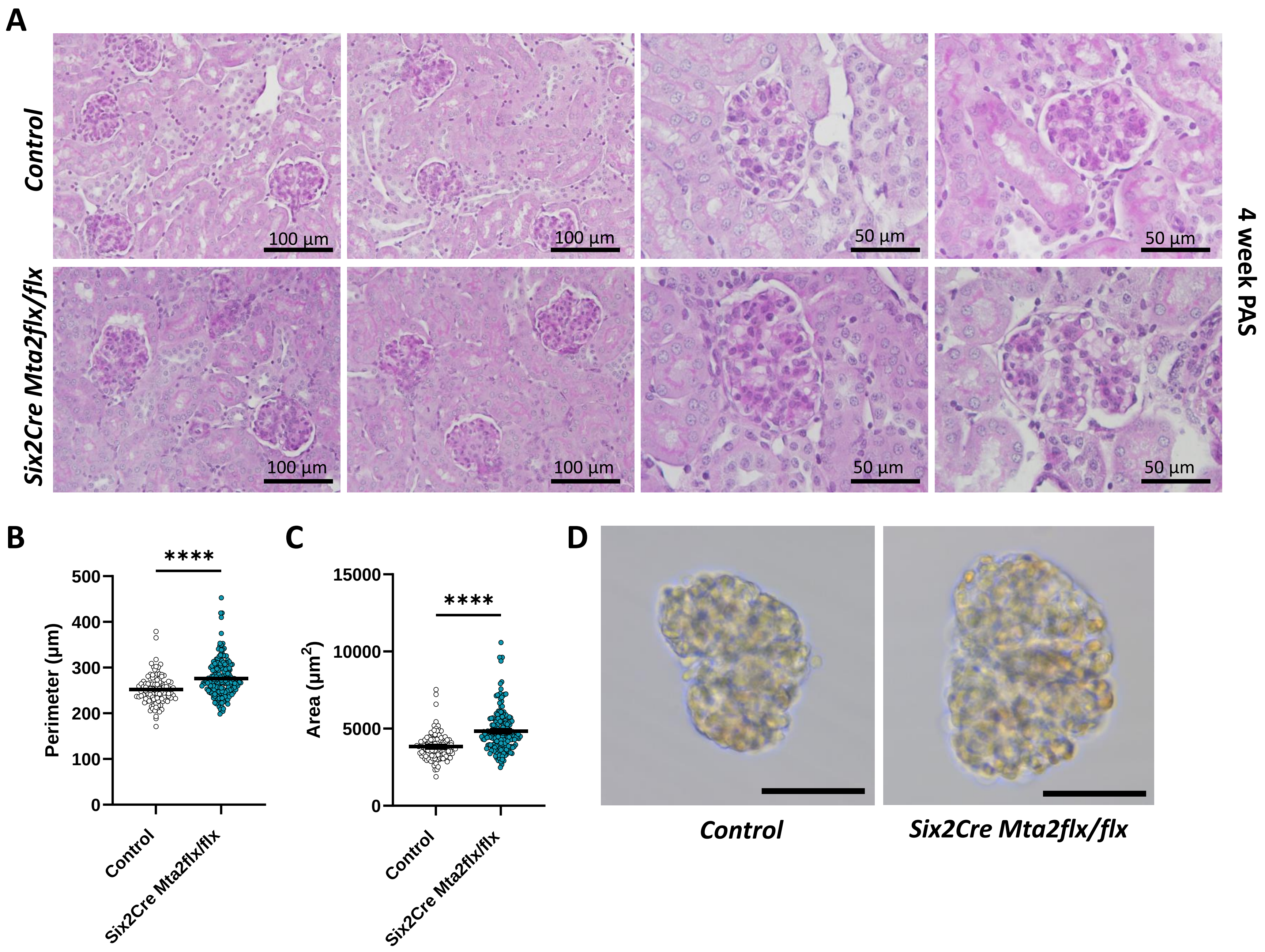
A
Control
Six2Cre Mta2flx/flx
100 µm
100 µm
50 µm
50 µm
100 µm
100 µm
50 µm
50 µm
4 week PAS
B
****
Perimeter (µm)
500
400
300
200
100
0
Control
Six2Cre Mta2flx/flx
C
****
Area (µm2)
15000
10000
5000
0
Control
Six2Cre Mta2flx/flx
D
Control
Six2Cre Mta2flx/flx

# S8

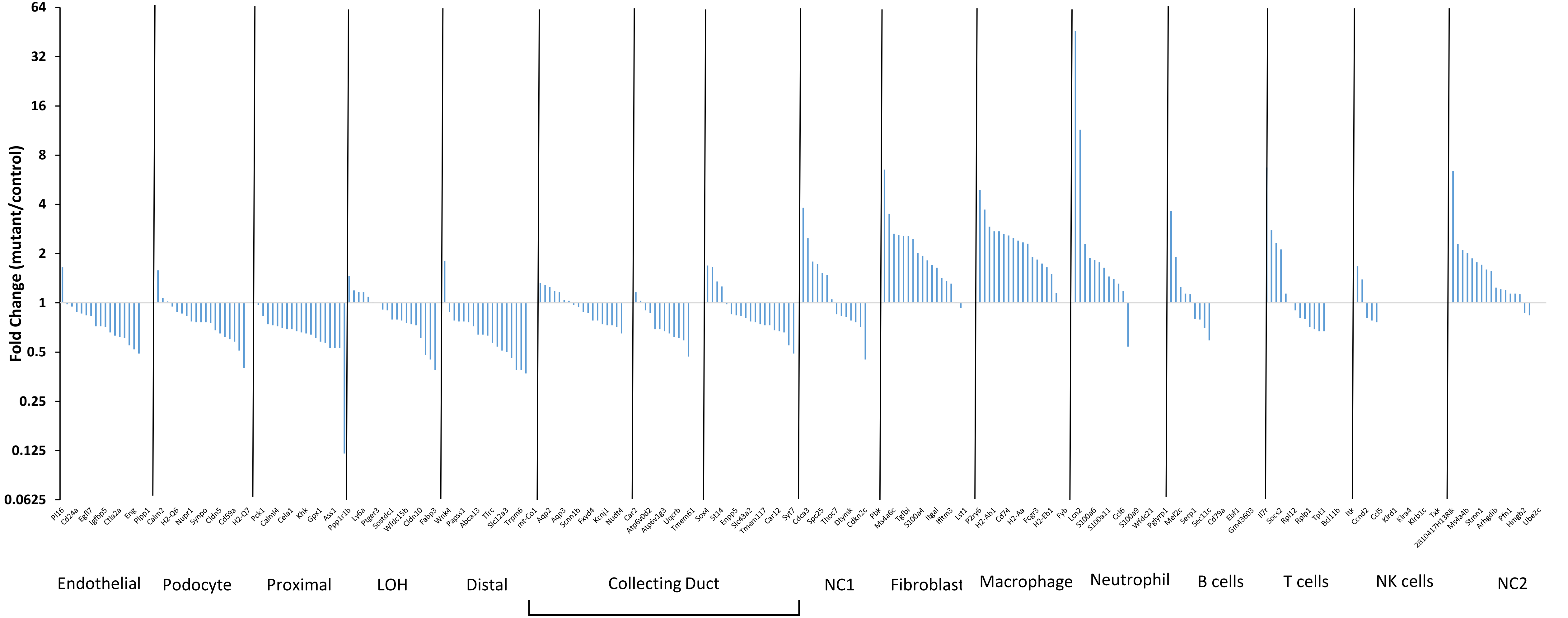
Fold Change (mutant/control)
64
32
16
8
4
2
1
0.5
0.25
0.125
0.0625
Pi16 Cd24a Egfl7 Igfbp5 Ctla2a Eng Plpp1
Calm2 H2-Q6 Nupr1 Synpo Cldn5 Cd59a H2-Q7
Pck1 Calml4 Cela1 Khk Gpx1 Ass1 Ppp1r1b
Ly6a Ptger3 Sostdc1 Wfdc15b Cldn10 Fabp3
Wnk4 Papss1 Abca13 Tfrc Slc12a3 Trpm6 mt-Co1
Aqp2 Aqp3 Scnn1b Fxyd4 Kcnj1 Nudt4
Car2 Atp6v0d2 Atp6v1g3 Uqcrb Tmem61
Sox4 St14 Enpp5 Slc43a2 Tmem117 Car12 Syt7
Cdca3 Spc25 Thoc7 Dtymk Cdkn2c
Pbk Ms4a6c Tgfbi S100a4 Itgal Ifitm3 Lst1
P2ry6 H2-Ab1 Cd74 H2-Aa Fcgr3 H2-Eb1 Fyb
Lcn2 S100a6 S100a11 Ccl6 S100a9 Wfdc21 Pglyrp1
Mef2c Serp1 Sec11c Cd79a Ebf1 Gm43603
Il7r Socs2 Rpl12 Rplp1 Tpt1 Bcl11b
Itk Ccnd2 Ccl5 Klrd1 Klra4 Klrb1c Txk
2810417H13Rik Ms4a4b Stmn1 Arhgdib Pfn1 Hmgb2 Ube2c
Endothelial
Podocyte
Proximal
LOH
Distal
Collecting Duct
NC1
Fibroblast
Macrophage
Neutrophil
B cells
T cells
NK cells
NC2

**S9**

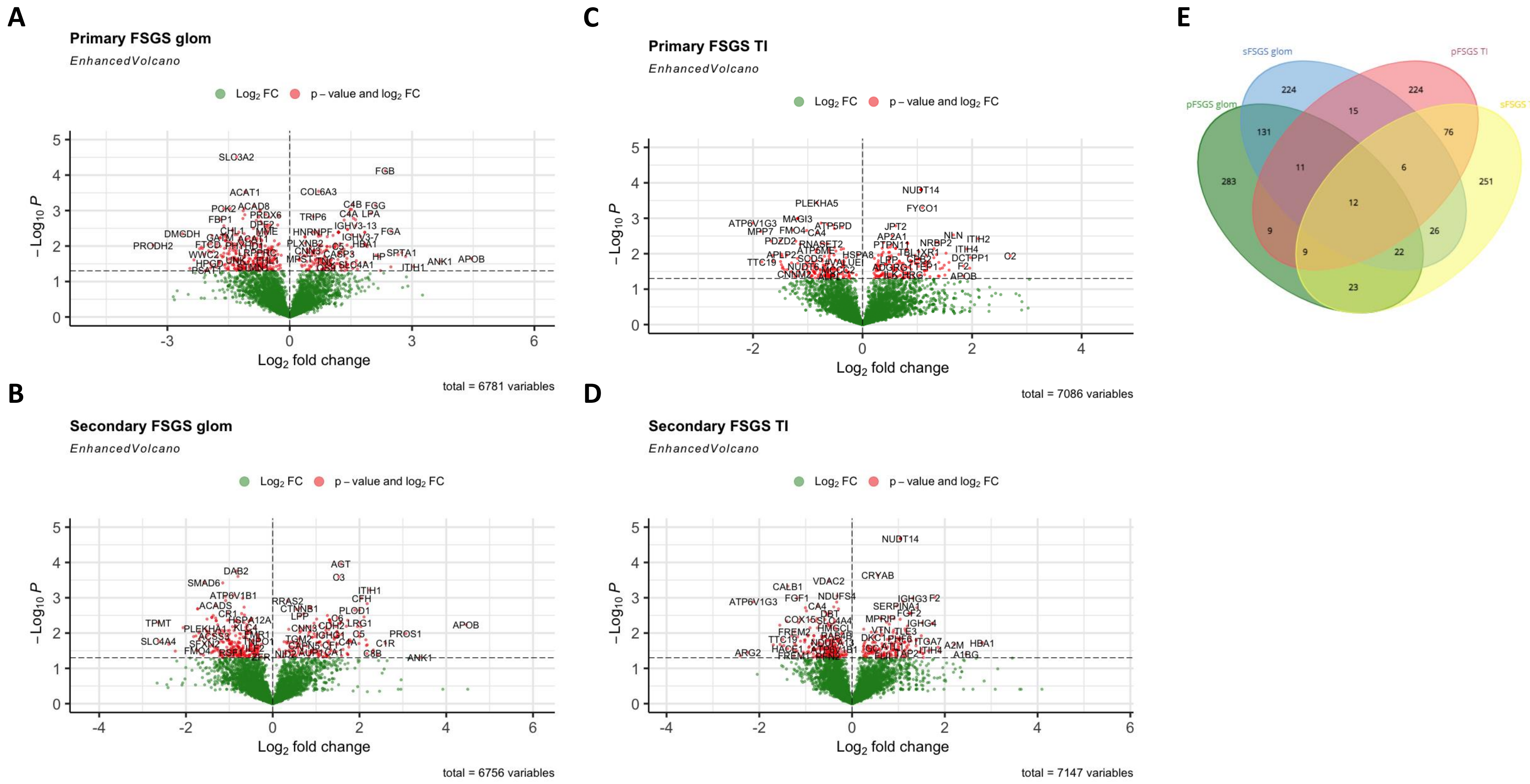
A
Primary FSGS glom
EnhancedVolcano
Log2 FC
p – value and log2 FC
−Log10 P
Log2 fold change
total = 6781 variables
B
Secondary FSGS glom
EnhancedVolcano
Log2 FC
p – value and log2 FC
−Log10 P
Log2 fold change
total = 6756 variables
C
Primary FSGS TI
EnhancedVolcano
Log2 FC
p – value and log2 FC
−Log10 P
Log2 fold change
total = 7086 variables
D
Secondary FSGS TI
EnhancedVolcano
Log2 FC
p – value and log2 FC
−Log10 P
Log2 fold change
total = 7147 variables
E
sFSGS glom
pFSGS TI
pFSGS glom
sFSGS TI
224
224
15
131
76
11
6
283
251
12
9
26
9
22
23

**Supp. Figure 1.** E17 *Mta2* RNA-seq compared to E18 single cell clusters. Fold change expression (mutant/control) values plotted for the top 20 DEGs from single cell RNA-seq cell clusters determined from E18 mouse kidney by the Melissa Little lab (*Development 2019 146: dev178673 doi: 10.1242/dev.178673*). Cell clusters for developing nephron structures (S-shaped body, early proximal, proximal, and podocyte) have a majority of genes in the *Mta2* mutant that are downregulated. For space, not all 20 DEGs are labeled.

**Supp. Figure 2.** Body weights for control and mutant male and female mice. Body weights (g) from birth (P0) to 1 year of age of control and *Six2Cre Mta2flx/flx* male and female mice. No significant differences were observed between control and mutant at any time point (multiple paired t-tests, Prism).

**Supp. Figure 3.** PAS staining of sections from 2 and 6 month control and *Mta2* mutant kidney. Glomeruli from these representative images were used to determine the semi-quantitative glomerulosclerosis score (Fig. 4D). The glomeruli were graded 0 (no change) to 4 (>75% of glomerulus PAS positive). Scale bar = 50 µm.

**Supp. Figure 4.** Masson's trichrome staining of sections from 2 and 6 month control and mutant *Mta2* kidney. Blue stain deposition highlighting collagen deposition in mutant 2 and 6 month kidney in the tubulointerstitial space and glomeruli. Left panels scale bar = 500 µm, right panels scale bar = 100 µm.

**Supp. Figure 5.** Hematoxylin and eosin staining of sections from 2 and 6 month control and mutant *Mta2* kidney. Tubular dilation, tubular damage, and glomerular damage is evident at 2 and 6 months in the mutant kidney. Left panels scale bar = 500 µm, right panels scale bar = 200 µm.

**Supp. Figure 6.** PAS staining of sections from 2 and 6 month control and *Mta2* mutant kidney. The HPF from these representative images were used to determine the semi-quantitative tubulointerstitial injury score (Fig. 4E). The HPF were graded 0 (no change) to 4 (<75% of area damaged). Loss of brushed border in proximal tubule, tubule dilation, and protein casts are evident in the mutant. Scale bar = 50 µm.

**Supp. Figure 7. A.** PAS staining of sections from 4-week-old control and mutant *Mta2* kidney. No PAS deposition was evident in the mutant kidney; however, the glomeruli in the mutant kidney were larger. 3 biological replicates were used for control and mutant. Left panels scale bar = 100 µm, right panels scale bar = 50 µm. **B.** Quantification of glomerular perimeter of 4-

week-old *Mta2flx/flx* control and *Six2Cre Mta2flx/flx* mutant glomeruli isolated using the differential adhesion method. Perimeter of mutant glomeruli were larger than the control glomeruli, Student's t-test, $p<0.0001$. **C.** Quantification of glomerular area of 4 week old control and mutant glomeruli isolated using the differential adhesion method. The area of mutant glomeruli were larger than glomeruli from control mice, Student's t-test, $p<0.001$. For **B, C,** 111 glomeruli were analyzed from 2 biological replicates for control and 197 glomeruli from 4 biological replicates for mutant. **D.** Representative images of glomeruli from control and mutant 4-week-old kidney isolated using the differential adhesion method used to quantify perimeter and area. Scale bar = 50 µm.

**Supp. Figure 8.** 2 month *Mta2* RNA-seq compared to adult mouse kidney single cell clusters. Fold change expression (mutant/control) values plotted for the top 20 DEGs from single cell RNA-seq cell clusters determined from adult mouse kidney by the Katalin Susztak Lab (*Science. 2018 May 18; 360(6390): 758–763*). Cell clusters for proximal nephron lineages (podocyte, proximal tubule) have downregulated expression in *Mta2* mutant kidney. Similarly, loop of Henle and distal tubule cell cluster genes also have downregulated expression in *Mta2* mutant kidney. Immune cell clusters (macrophage, neutrophil, T-cells) have upregulated gene expression in the *Mta2* mutant kidney. For space, not all 20 DEGs are labeled.

**Supp. Figure 9.** Human biopsy regional proteomics. **A-D.** Volcano plot of differentially expressed proteins ($p<0.05$, red) in primary and secondary FSGS glomeruli (glom), and primary and secondary FSGS tubulointerstitia (TI) compared with normal kidney. **E.** Venn diagram displaying the overlap of differentially expressed proteins ($p<0.05$) between primary and secondary FSGS glomeruli (glom) and tubulointerstitia (TI).
